# Rapid evolution of the human mutation spectrum

**DOI:** 10.1101/084343

**Authors:** Kelley Harris, Jonathan K. Pritchard

## Abstract

DNA is a remarkably precise medium for copying and storing biological information. This high fidelity results from the action of hundreds of genes involved in replication, proofreading, and damage repair. Evolutionary theory suggests that in such a system, selection has limited ability to remove genetic variants that change mutation rates by small amounts or in specific sequence contexts. Consistent with this, using SNV variation as a proxy for mutational input, we report here that mutational spectra differ substantially among species, human continental groups and even some closely-related populations. Close examination of one signal, an increased TCC*→*TTC mutation rate in Europeans, indicates a burst of mutations from about 15,000 to 2,000 years ago, perhaps due to the appearance, drift, and ultimate elimination of a genetic modifier of mutation rate. Our results suggest that mutation rates can evolve markedly over short evolutionary timescales and suggest the possibility of mapping mutational modifiers.

## Main Text

Germline mutations provide the raw material for evolution, but also generate genetic load and inherited disease. Indeed, the vast majority of mutations that affect fitness are deleterious, and hence biological systems have evolved elaborate mechanisms for accurate DNA replication and repair of diverse types of spontaneous damage. Due to the combined action of hundreds of genes, mutation rates are extremely low–in humans, about 1 point mutation per 100MB or about 60 genome-wide per generation [1, 2].

While the precise roles of most of the relevant genes have not been fully elucidated, research on somatic mutations in cancer has shown that defects in particular genes can lead to increased mutation rates within very specific sequence contexts [3, 4]. For example, mutations in the proofreading exonuclease domain of DNA polymerase ϵ cause TCT*→*TAT and TCG*→*TTG mutations on the leading DNA strand [5]. Mutational shifts of this kind have been referred to as “mutational signatures”. Specific signatures may also be caused by nongenetic factors such as chemical mutagens, UV damage, or guanine oxidation [6].

Together, these observations imply a high degree of specialization of individual genes involved in DNA proofreading and repair. While the repair system has evolved to be extremely accurate overall, theory suggests that in such a system, natural selection may have limited ability to fine-tune the efficacy of individual genes [7, 8]. If a variant in a repair gene increases or decreases the overall mutation rate by a small amount–for example, only in a very specific sequence context–then the net effect on fitness may fall below the threshold at which natural selection is effective. (Drift tends to dominate selection when the change in fitness is less than the inverse of effective population size). The limits of selection on mutation rate modifiers are especially acute in recombining organisms such as humans because a variant that increases the mutation rate can recombine away from deleterious mutations it generates elsewhere in the genome.

Given these theoretical predictions, we hypothesized that there may be substantial scope for modifiers of mutation rates to segregate within human populations, or between closely related species. Most triplet sequence contexts have mutation rates that vary across the evolutionary tree of mammals [9], but evolution of the mutation spectrum over short time scales has been less well described. Weak natural mutators have recently been observed in yeast [10] and inferred from human haplotype data [11]; if such mutators affect specific pathways of proofreading or repair, then we may expect shifts in the abundance of mutations within particular sequence contexts. Indeed, one of us has recently identified a candidate signal of this type, namely an increase in TCC*→*TTC transitions in Europeans, relative to other populations [12]; this was recently replicated [13]. Here we show that mutation spectrum change is much more widespread than these initial studies suggested: although the TCC*→*TTC rate increase in Europeans was unusually dramatic, smaller-scale changes are so commonplace that almost every great ape species and human continental group has its own distinctive mutational spectrum.

## Results

To investigate the mutational processes in different human populations, we classified each single nucleotide variants (SNV) in the 1000 Genomes Phase 3 data [14] in terms of its ancestral allele, derived allele, and 5’ and 3’ flanking nucleotides. We collapsed strand complements together to obtain 96 SNV categories. Since the detection of singletons may vary across samples, and because some singletons may result from cell-line or somatic mutations, we only considered variants seen in more than one copy. We further excluded variants in annotated repeats (since read mapping error rates may be higher in such regions) and in PhyloP conserved regions (to avoid selectively constrained regions) [15]. From the remaining sites, we calculated the distribution of derived SNVs carried by each Phase 3 individual. We used this as a proxy for the mutational input spectrum in the ancestors of each individual.

To explore global patterns of the mutation spectrum, we performed principal component analysis (PCA) in which each individual was characterized simply by the fraction of their derived alleles in each of the 96 SNV categories (Fig. 1A). PCA is commonly applied to individual-level genotypes, in which case the PCs are usually highly correlated with geography [16]. Although the triplet mutation spectrum is an extremely compressed summary statistic compared to typical genotype arrays, we found that it contains sufficient information to reliably classify individuals by continent of origin. The first principal component separated Africans from non-Africans, and the second separated Europeans from East Asians, with South Asians and admixed native Americans (Figure 1–Figure Supplement 2) appearing intermediate between the two.

**Figure 1:**
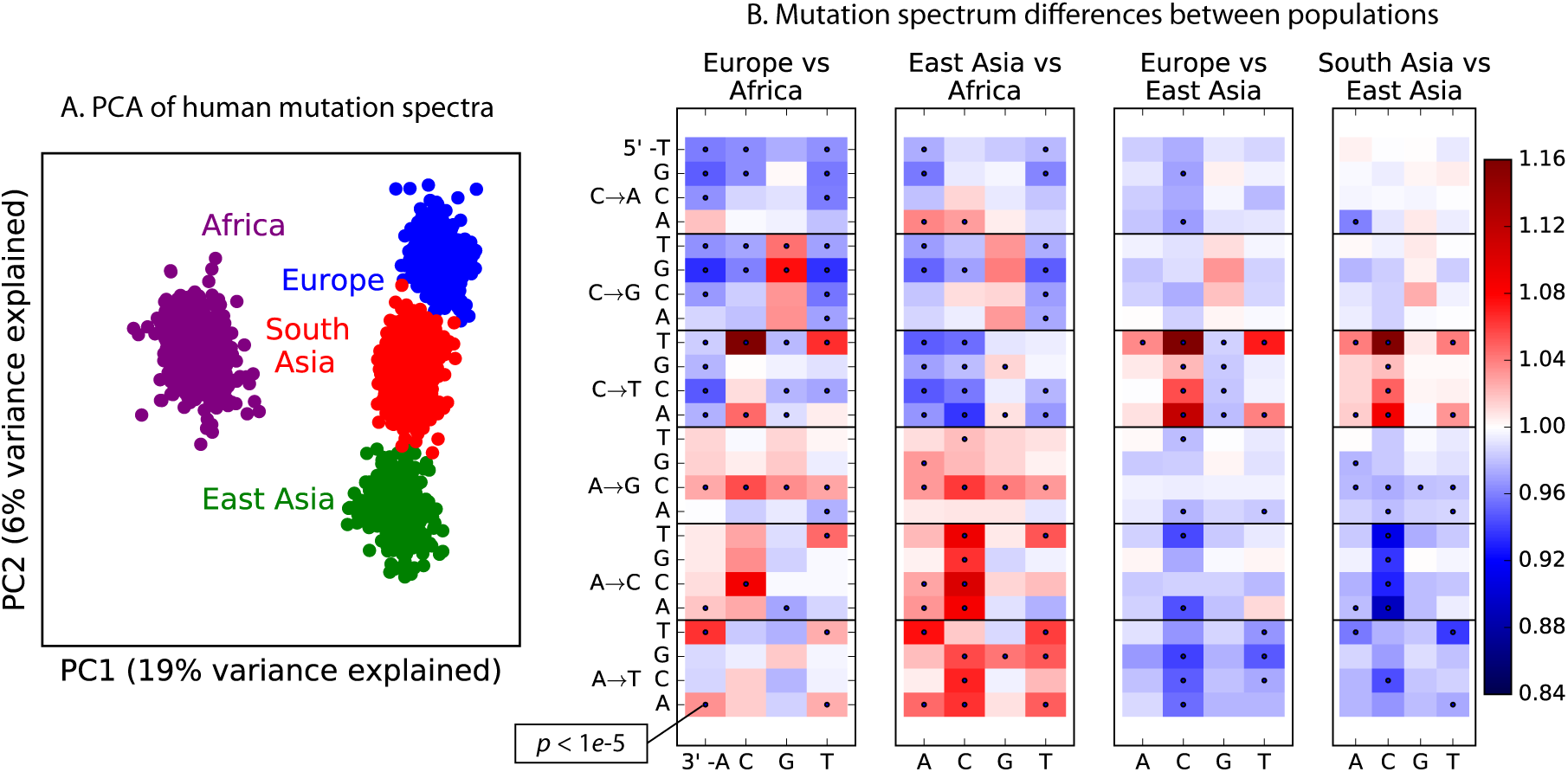
Global patterns of variation in SNV spectra. A. *Principal Component Analysis of individuals according to the fraction of derived alleles that each individual carries in each of 96 mutational types.* **B.** *Heatmaps showing, for pairs of continental groups, the ratio of the proportions of SNVs in each of the 96 mutational types. Each block corresponds to one mutation type; within blocks, rows indicate the 5’ nucleotide, and columns indicate the 3’ nucleotide. Red colors indicate a greater fraction of a given mutation type in the first-listed group relative to the second. Points indicate significant contrasts at p <* 10*−*^5^*. See Figure Supplements 1, 2, and 3 for heatmap comparisons between additional population pairs as well as a description of PCA loadings and the p-values of all mutation class enrichments. Figure Supplement 4 demonstrates that these patterns are unlikely to be driven by biased gene conversion. In Figure Supplement 5, we see that this mutation spectrum structure replicates on both strands of the transcribed genome as well as the non-transcribed portion of the genome. Figure Supplements 6, 7, and 8 show that most of this structure replicates across multiple chromatin states and varies little with replication timing.*

Remarkably, we found that the mutation spectrum differences among continental groups are composed of small shifts in the abundance of many different mutation types (Fig. 1B). For example, comparing Africans and Europeans, 43 of the 96 mutation types are significant at a *p <* 10*−*^5^ threshold using a forward variable selection procedure. The previously described TCC*→*TTC signature partially drives the difference between Europeans and the other groups, but most other shifts are smaller in magnitude and appear to be spread over more diffuse sets of related mutation types. East Asians have excess A*→*T transversions in most sequence contexts, as well as about 10% more *AC*→*\*CC mutations than any other group. Compared to Africans, all Eurasians have proportionally fewer C*→*\* mutations relative to A*→*\* mutations.

### Replication of mutation spectrum shifts

One possible concern is that batch effects or other sequencing artifacts might contribute to differences in mutational spectra. Therefore we replicated our analysis using 201 genomes from the Simons Genome Diversity Project [17]. The SGDP genomes were sequenced at high coverage, independently from 1000 Genomes, using an almost non-overlapping panel of samples. We found extremely strong agreement between the mutational shifts in the two data sets (Fig. 2). For example, all of the 43 mutation types with a significant difference between Africa and Europe (at *p <* 10^−5^) in 1000 Genomes also show a frequency difference in the same direction in SGDP (comparing Africa and West Eurasia). In both 1000 Genomes and SGDP, the enrichment of *AC*→*\*CC mutations in East Asia is larger in magnitude than any other signal aside from the previously described TCC*→*TTC imbalance.

**Figure 2:**
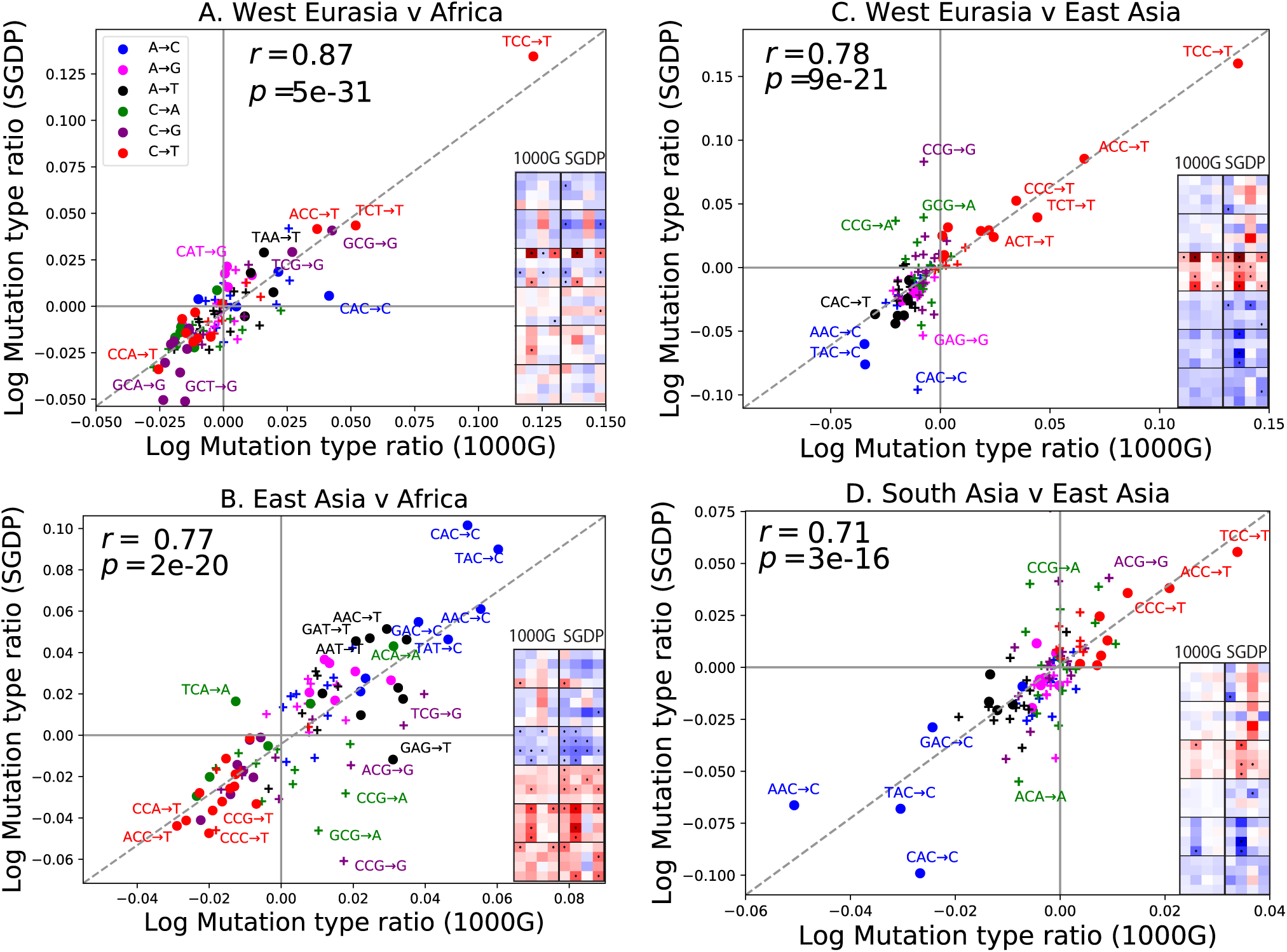
Concordance of mutational shifts in 1000 Genomes vs. SGDP. *Each panel shows natural-log mutation spectrum ratios between a pair of continental groups, based on 1000 Genomes (x-axis) and SGDP (y-axis) data. Data points encoded by (+) symbols denote mutation types that are not significantly enriched in either population in the Figure 1 1000 Genomes analysis (p <* 10^−5^*). These heatmaps use the same labeling and color scale as in Figure 1. All 1000 Genomes ratios in this figure were estimated after projecting the 1000 Genomes site frequency spectrum down to the sample size of SGDP. See Figure Supplements 1 and 2 for a complete set of SGDP heatmaps and regressions versus 1000 Genomes.*

The greatest discrepancies between 1000 Genomes and SGDP involve transversions at CpG sites, which are among the rarest mutational classes. These discrepancies might result from data processing differences or random sampling variation, but might also reflect differences in the fine-scale ethnic composition of the two panels.

### Evidence for a pulse of TCC*→*TTC mutations in Europe and South Asia

To investigate the timescale over which the mutation spectrum change occurred, we analyzed the allele frequency distribution of TCC*→*TTC mutations, which are highly enriched in Europeans (Fig. 3A; *p <* 1 *×* 10^−300^ for Europe vs. Africa) and to a lesser extent in South Asians. We calculated allele frequencies both in 1000 Genomes and in the larger UK10K genome panel [18]. As expected for a signal that is primarily European, we found particular enrichment of these mutations at low frequencies. But surprisingly, the enrichment peaks around 0.6% frequency in UK10K, and there is practically no enrichment among the very lowest frequency variants (Figure 3B and Figure 3–Figure Supplement 1). C*→*T mutations on other backgrounds, namely within TCT, CCC and ACC contexts, are also enriched in Europe and South Asia and show a similar enrichment around 0.6% frequency that declines among rarer variants (Fig. 3C). This suggests that these four mutation types comprise the signature of a single mutational pulse that is no longer active. No other mutation types show such a pulse-like distribution in UK10K, though several types show evidence of monotonic rate change over time (Figure 3–Figure Supplements 3,4 and 5).

**Figure 3:**
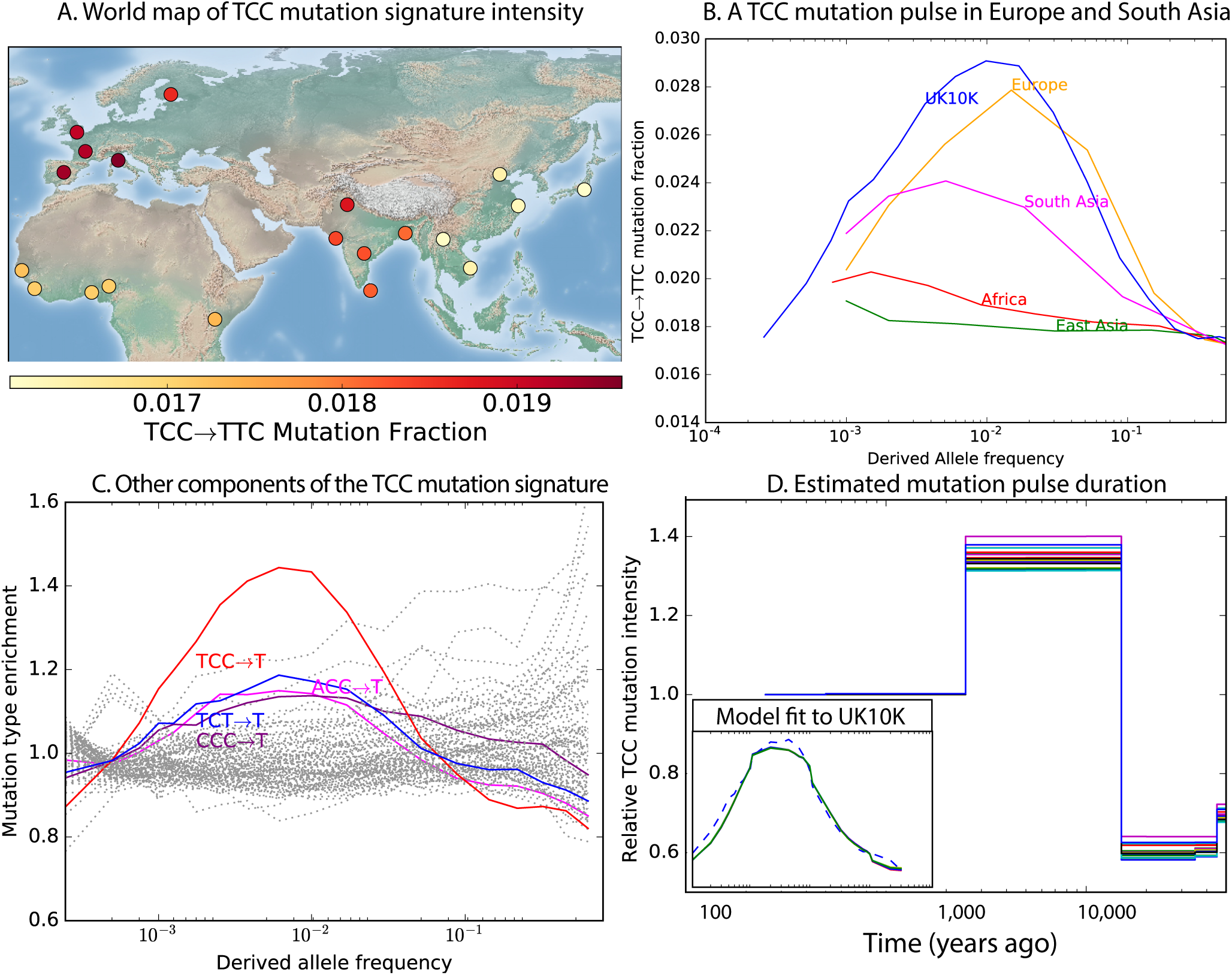
Geographic distribution and age of the TCC mutation pulse. **(A)** *Observed frequencies of TCC TTC variants in 1000 Genomes populations.* **(B)** *Fraction of TCC TTC variants as a function of allele frequency in different samples indicates that these peak around 1%. See Figure Supplement 1 for distributions of TCC TTC allele frequency within all 1000 Genomes populations, and see Figure Supplement 2 for the replication of this result in the Exome Aggregation Consortium Data. In the UK10K data, which has the largest sample size, the peak occurs at 0.6% allele frequency.* **(C)** *Other enriched C T mutations with similar context also peak at 0.6% frequency in UK10K. See Figure Supplements 3, 4 and 5 for labeled allele frequency distributions of all 96 mutation types (most represented here as unlabeled grey lines). See Figure Supplement 6 for heatmap comparisons of the 1000 Genomes populations partitioned by allele frequency, which provide a different view of these patterns.* **(D)** *A population genetic model supports a pulse of TCC TTC mutations from 15,000–2,000 years ago. Inset shows the observed and predicted frequency distributions of this mutation under the inferred model.*

We used the enrichment of TCC*→*TTC mutations as a function of allele frequency to estimate when this mutation pulse was active. Assuming a simple piecewise-constant model, we infer that the rate of TCC*→*TTC mutations increased dramatically ~15,000 years ago and decreased again ~2,000 years ago. This time-range is consistent with results showing this signal in a pair of prehistoric European samples from 7,000 and 8,000 years ago, respectively [13]. We hypothesize that this mutation pulse may have been caused by a mutator allele that drifted up in frequency starting 15,000 years ago, but that is now rare or absent from present day populations.

Although low frequency allele calls often contain a higher proportion of base calling errors than higher frequency allele calls do, it is not plausible that base-calling errors could be responsible for the pulse we have described. In the UK10K data, a minor allele present at 0.6% frequency corresponds to a derived allele that is present in 23 out of 3854 sampled haplotypes and supported by 80 short reads on average (assuming 7x coverage per individual). When independently generated datasets of different sizes are projected down to the same sample size, the TCC*→*TTC pulse spans the same range of allele frequencies in both datasets (Figure 3–Figure Supplements 1 and 2), which would not be the case if the shape of the curve were a function of low frequency errors.

### Fine-scale mutation spectrum variation in other populations

Encouraged by these results, we sought to find other signatures of recent mutation pulses. We generated heatmaps and PCA plots of mutation spectrum variation within each continental group, looking for fine-scale differences between closely related populations (Figure 4 and Figure 4–Figure Supplements 1 through 6). In some cases mutational spectra differ even between very closely related populations. For example, the *AC*→*\*CC mutations with elevated rates in East Asia appear to be distributed heterogeneously within that group, with most of the load carried by a subset of the Japanese individuals. These individuals also have elevated rates of ACA*→*AAA and TAT*→*TTT mutations (Figure 4A and Figure Supplement 4). This signature appears to be present in only a handful of Chinese individuals, and no Kinh or Dai individuals. As seen for the European TCC mutation, the enrichment of these mutation types peaks at low frequencies, i.e., ~1%. Given the availability of only 200 Japanese individuals in 1000 Genomes, it is hard to say whether the true peak is at a frequency much lower than 1%.

**Figure 4:**
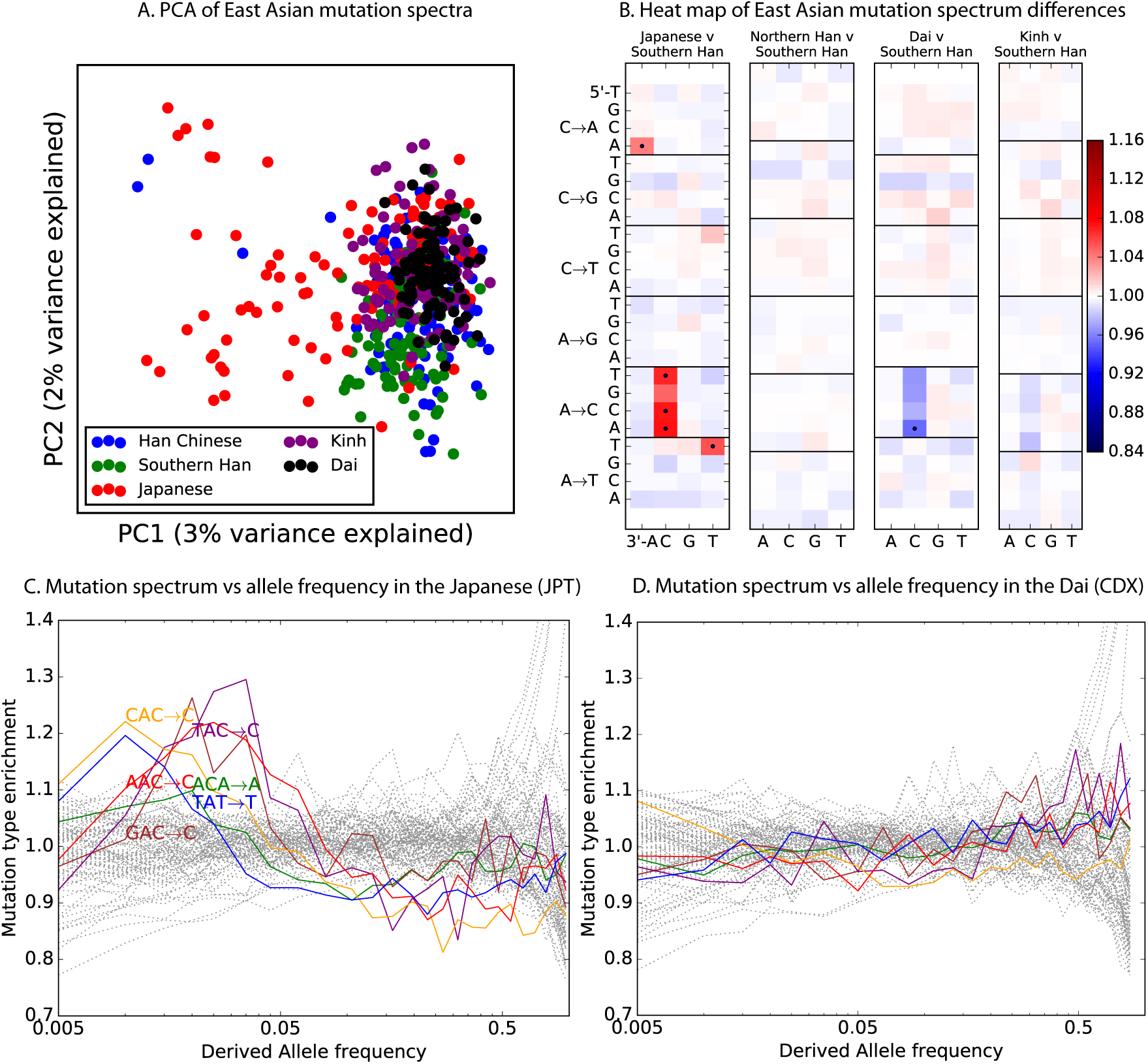
Mutational variation among east Asian populations. **(A)** *PCA of east Asian samples from 1000 Genomes, based on the relative proportions of each of the 96 mutational types. See Figure Supplements 2 through 6 for other finescale population PCAs.* **(B)** *Heatmaps showing, for pairs of east Asian samples, the ratio of the proportions of SNVs in each of the 96 mutational types. Points indicate significant contrasts at p <* 10*−*^5^*. See Figure Supplement 1 for additional finescale heatmaps.* **(C)** *and* **(D)** *Relative enrichment of each mutational type in Japanese and Dai, respectively as a function of allele frequency. Six mutation types that are enriched in JPT are indicated. Populations: CDX=Dai, CHB=Han (Beijing); CHS=Han (south China); KHV=Kinh; JPT=Japanese.*

PCA reveals relatively little fine-scale structure within the mutational spectra of Europeans or South Asians (Figure 4–Figure Supplements 5 and 6). However, Africans exhibit some substructure (Figure 4–Figure Supplement 3), with the Luhya exhibiting the most distinctive mutational spectrum. Unexpectedly, a closer examination of PC loadings reveals that the Luhya outliers are enriched for the same mutational signature identified in the Japanese. Even in Europeans and South Asians, the first PC is heavily weighted toward *AC*→*\*CC, ACA*→*AAA, and TAT*→*TTT, although this signature explains less of the mutation spectrum variance within these more homogeneous populations. The sharing of this signature may suggest either parallel increases of a shared mutation modifier, or a shared aspect of environment or life history that affects the mutation spectrum.

### Mutation spectrum variation among the great apes

Finally, given our finding of extensive fine-scale variation in mutational spectra between human populations, we hypothesized that mutational variation between species is likely to be even greater. To compare the mutation spectra of the great apes in more detail, we obtained SNV data from the Great Ape Diversity Panel, which includes 78 whole genome sequences from six great ape species including human [19]. Overall, we find dramatic variation in mutational spectra among the great ape species (Figure 5 and Figure 5–Figure Supplement 1).

As noted previously [20], one major trend is a higher proportion of CpG mutations among the species closest to human, possibly reflecting lengthening generation time along the human lineage, consistent with previous indications that species closely related to humans have lower mutation rates than more distant species [21, 22, 23]. However, most other differences are not obviously related to known processes such as biased gene conversion and generation time change. The A*→*T mutation rate appears to have sped up in the common ancestor of humans, chimpanzees, and bonobos, a change that appears consistent with a mutator variant that was fixed instead of lost. It is unclear whether this ancient A*→*T speedup is related to the A*→*T speedup in East Asians. Other mutational signatures appear on only a single branch of the great ape tree, such as a slowdown of A*→*C mutations in gorillas.

## Discussion

The widespread differences captured in Figures 1 and 2 may be footprints of allele frequency shifts affecting different mutator alleles. But in principle, other genetic and non-genetic processes may also impact the observed mutational spectrum. First, biased gene conversion (BGC) tends to favor C/G alleles over A/T, and BGC is potentially more efficient in populations of large effective size compared to populations of smaller effective size [24]. However, despite the bottlenecks that are known to have affected Eurasian diversity, there is no clear trend of an increased fraction of C/G*→*A/T relative to A/T*→*C/G in non-Africans vs. Africans, or with distance from Africa (Figure 1–Figure Supplement 7), and previous studies have also found little evidence for a strong genome-wide effect of BGC on the mutational spectrum in humans and great apes [26, 20]. For these reasons, we think that evolution of the mutational process is a better explanation than BGC or selection for differences that have been observed between the spectra of ultra-rare singleton variants and older human genetic variation [25];

It is also known that shifts in generation time or other life-history traits may affect mutational spectra, particularly for CpG transitions [27, 28]. Most CpG transitions result from spontaneous methyl-cytosine deamination as opposed to errors in DNA replication. Hence the rate of CpG transitions is less affected by generation time than other mutations [9, 29, 30]. We observe that Europeans have a lower fraction of CpG variants compared to Africans, East Asians and South Asians (Fig. 1B), consistent with a recent report of a lower rate of *de novo* CCG*→*CTG mutations in European individuals compared to Pakistanis [31]. Such a pattern may be consistent with a shorter average generation time in Europeans [29], though it is unclear that a plausible shift in generation-time could produce such a large effect. Apart from this, the other patterns evident in Figure 1 do not seem explainable by known processes.

In summary, we report here that, mutational spectra differ significantly among closely related human populations, and that they differ greatly among the great ape species. Our work shows that subtle, concerted shifts in the frequencies of many different mutation types are more widespread than dramatic jumps in the rate of single mutation types, although the existence of the European TCC*→*TTC pulse shows that both modes of evolution do occur [12, 29, 13].

At this time, we cannot exclude a role for nongenetic factors such as changes in life history or mutagen exposure in driving these signals. However given the sheer diversity of the effects reported here, it seems parsimonious to us to propose that most of this variation is driven by the appearance and drift of genetic modifiers of mutation rate. This situation is perhaps reminiscent of the earlier observation that genome-wide recombination patterns are variable among individuals [32], and ultimate discovery of PRDM9 [33]; although in this case it is unlikely that a single gene is responsible for all signals seen here. As large datasets of *de novo* mutations become available, it should be possible to map mutator loci genome-wide. In summary, our results suggest the likelihood that mutational modifiers are an important part of the landscape of human genetic variation.

## Acknowledgements

This work was funded by NIH grants GM116381 and HG008140, and by the Howard Hughes Medical Institute. We thank Jeffrey Spence and Yun S. Song for technical assistance. We also thank Ziyue Gao, Arbel Harpak, Molly Przeworski, Joshua Schraiber, and Aylwyn Scally for comments and discussion, as well as two anonymous reviewers.

## Methods

### Data Availability

All datasets analyzed here are publicly available at the following websites:

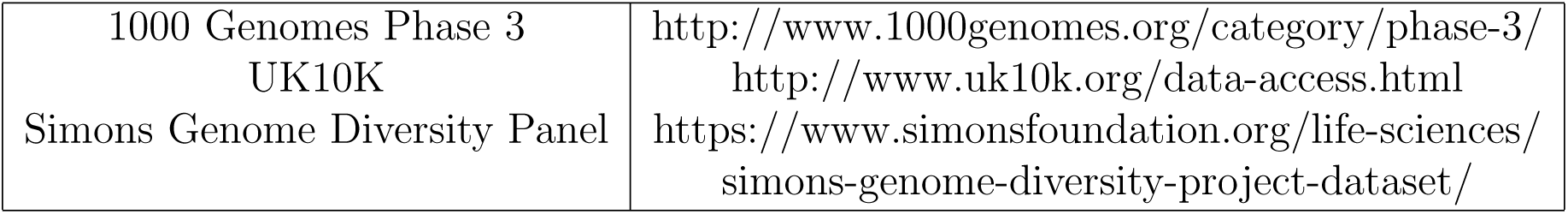

### Human Mutation Spectrum Processing

Mutation spectra were computed using 1000 Genomes Phase 3 SNPs [14] that are biallelic, pass all 1000 Genomes quality filters, and are not adjacent to any N’s in the hg19 reference sequence. Ancestral states were assigned using the UCSC Genome Browser alignment of hg19 to the PanTro2 chimpanzee reference genome; SNPs were discarded if neither the reference nor alternate allele matched the chimpanzee reference. To minimize the potential impact of ancestral misidentification errors, SNPs with derived allele frequency higher than 0.98 were discarded. We also filtered out regions annotated as “conserved” based on the 100-way PhyloP conservation score [15], download from http://hgdownload.cse.ucsc.edu/goldenPath/hg19/phastCons100way/, as well as regions annotated as repeats by RepeatMasker [34], downloaded from http://hgdownload.cse.ucsc.edu/goldenpath/hg19/database/nestedRepeats.txt.gz. To be counted as part of the mutation spectrum of population *P* (which can be either a continental group or a finer-scale population from one city), a SNP should not be a singleton within population *P*–at least two copies of the ancestral and derived alleles must be present within that population.

An identical approach was used to extract the mutation spectrum of the UK10K ALSPAC panel [18], which is not subdivided into smaller populations. The data were filtered as described in [35]. The filtering procedure performed by Field, et al. reduces the ALSPAC sample size to 1927 individuals.

We also computed mutation spectra of the Simons Genome Diversity Panel (SGDP) populations [17]. Four of the SGDP populations, West Eurasia, East Asia, South Asia, and Africa, were compared to their direct counterparts in the 1000 Genomes data. Three additional SGDP populations, Central Asia and Siberia, Oceania, and America, had no close 1000 Genomes counterparts and were not analyzed here (although each project contained a panel of people from the Americans, the composition of the American panels was extremely different, with the 1000 Genomes populations being much more admixed with Europeans and Africans). SGDP sites with more than 20% missing data were not utilized. All other data processing was done the same way described for the 1000 Genomes data.

The following table gives the same size of each population panel, as well as the total number of SNPs segregating in the panel that are used to compute mutation type ratios:

| Dataset | Population | Number of individuals | Number of SNPs |
| --- | --- | --- | --- |
| 1kg | Africa | 504 | 16,870,400 |
| 1kg | Europe | 503 | 8,508,040 |
| 1kg | East Asia | 504 | 7,895,925 |
| 1kg | South Asia | 489 | 9,552,781 |
| SGDP | Africa | 45 | 6,569,658 |
| SGDP | West Eurasia | 69 | 4,201,571 |
| SGDP | East Asia | 49 | 3,312,645 |
| SGDP | South Asia | 38 | 3,449,624 |

### Great Ape Diversity Panel Data Processing

Biallelic great ape SNPs were extracted from the Great Ape Diversity Panel VCF [19], which is aligned to the hg18 human reference sequence. Ancestral states were assigned using the Great Ape Genetic Diversity project annotation, which used the Felsenstein pruning algorithm to assign allelic states to internal nodes in the great ape tree. In the Great Ape Diversity Panel, the most recent common ancestor (MRCA) of the human species is labeled as node 18; the MRCAs of chimpanzees, bonobos, gorillas, and orangutans, respectively, are labeled as node 16, node 17, node 19, and node 15. We extracted the state of each MRCA at each SNP in the alignment and used it to polarize the ancestral and derived allele at that site; a SNP was discarded whenever the ancestral node was assigned an uncertain or polymorphic ancestral state. As with the human data, SNPs with derived allele frequency higher than 0.98 were not used, and both repeats and PhyloP-annotated conserved regions were filtered away.

### Visual representation of mutation spectra

The mutation type of a SNP is defined in terms of its ancestral allele, its derived allele, and its two immediate 5’ and 3’ neighbors. Two mutation types are considered equivalent if they are strand-complementary to each other (e.g. ACG*→*ATG is equivalent to CGT*→*CAT). This scheme classifies SNPs into 96 different mutation types, each that can be represented with an A or C ancestral allele.

To compute the frequency *fP* (*m*) of SNP *m* in population *P*, we count up all SNPs of type *m* where the derived allele is present in at least one representative of population *P* (which can be either a specific population such as YRI or a broader continental group such as AFR). After obtaining this count *CP* (*m*), we define *fP* (*m*) to be the ratio *CP* (*m*)*/ ∑*_*m’*_ *CP* (*m’*), where the sum in the denominator ranges over all 96 mutation types *m’*. The enrichment of mutation type *m* in population *P*_1_ relative to population *P*_2_ is defined to be *fP*_1_ (*m*)*/fP*_2_ (*m*); these enrichments are visualized as heat maps in Figures 1B, 3B, and 4A.

To track changes in the mutational spectrum over time, we compute *f*_*P*_ (*m*) in bins of restricted allele frequency. This involves counting the number of SNPs of type *m* that are present at frequency *ϕ* in population *P* to obtain counts *C*_*P*_ (*m, ϕ*) and frequencies *fP* (*m, ϕ*) = *CP* (*m, ϕ*)*Σ_m’_ CP* (*m’ϕ*). Deviation of the ratio *fP* (*m, ϕ*)*/fP* (*m*) from 1 indicates that the rate of *m* has fluctuated recently in the history of population *P*. To make the sampling noise approximately uniform across alleles of different frequencies, alleles of derived count greater than 5 were grouped into approximately log-spaced bins that each contained similar numbers of UK10K SNPs. More precisely, we defined a set of bin endpoints *b*_1_*, b*_2_, … such that the total number of SNPs ranging in derived allele count between *b_i_* and *b*_*i*_+1 *−* 1 is greater than or equal to the number of 5-ton SNPs, while the total number of SNPs ranging in derived allele count from *b_i_* to *b*_*i*_+1 *−* 2 is less than the number of 5-ton SNPs.

In some cases, e.g. Figures 2, Figure 2–Figure Supplement 1B, and Figure 3–Figure Supplement 1, site frequency spectra were projected down to a smaller sample size before counting SNPs in order to more accurately compare datasets of different sample sizes. A binomial sampling approach was used to project a sample of *N* haplotypes does to a smaller sample size *n*. Letting 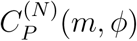 denote the SNP counts in the large sample of *N* haplotypes, effective SNP counts 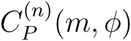 in a sample of *n* haplotypes are computed as follows:

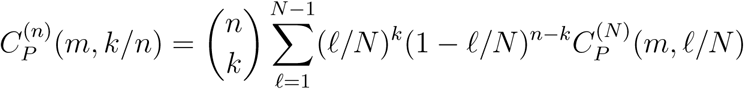

### Significance testing

One central goal of this paper is to test whether many mutation types differ in rate between human populations or whether mutation spectrum shifts have been rare events affecting only a small proportion of mutation types. A simple statistical method for answering this question would be to perform 96 separate chi-square tests, one for each triplet-context-dependent mutation type, as follows:

Let *S_i_* denote the total number of SNPs segregating in population *P_i_*, and let 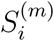 denote the number of SNPs of mutation type *m*. If mutation type *m* is more prevalent in population *P*_1_ than in population *P*_2_, a chi-square test provides a natural way of assessing the significance of this difference. As described in [12], this test is performed on the following 2-by-2 contingency table:

|  |  |
| --- | --- |
| $S_1^{(m)}$ | $P_1 - S_1^{(m)}$ |
| $S_2^{(m)}$ | $P_2 - S_2^{(m)}$ |

It would be appealing to conclude that every mutation type “passing” this chi-square test is a mutation type that has changed in rate during recent human history. However, if we were to perform the full set of 96 tests, they would not be independent. A sufficiently large increase in the rate of one mutation type *m*_1_ in population *P*_1_ after divergence from *P*_2_ could cause another mutation type *m*_2_, whose rate has remained constant, to comprise significantly different fractions of the SNPs from *P*_1_ and *P*_2_. To minimize this effect, we formulate the following iterative procedure of conditionally independent tests: first, compute a chi-square significance value *p*_unordered_(*m*) for each mutation type *m* using the 2-by-2 chi-square table above. We then use these values to order the SNPs from lowest *p* value to highest and compute a set of ordered *p* values *p*_ordered_(*m*). For the mutation type *m*_0_ with the lowest unordered *p* value, *p*_unordered_(*m*_0_) = *p*_ordered_(*m*_0_). For mutation type *m*_*i*_, which has the *i*th lowest unordered *p* value and *i <* 96, *p*_ordered_(*m_i_*) is computed from the following contingency table:

|  |  |
| --- | --- |
| $S_1^{(m_i)}$ | $\sum_{j=i+1}^{96} S_1^{(m_j)}$ |
| $S_2^{(m_i)}$ | $\sum_{j=i+1}^{96} S_2^{(m_j)}$ |

For mutation type *m*_96_, which has the highest unordered *p* value, the ordered *p* value is computed from the contingency table

|  |  |
| --- | --- |
| $S_1^{(m_{96})}$ | $S_1^{(m_{95})}$ |
| $S_2^{(m_{96})}$ | $S_2^{(m_{95})}$ |

This procedure is guaranteed to find fewer mutation types to differ significantly in rate between populations compared to separate chi-square tests.

### Principal component analysis (PCA)

The python package matplotlib.mlab.PCA was used to perform PCA on the complete set of 1000 Genomes haplotypes, each haplotype *h* represented by a 96-element vector encoding the mutation frequencies (*f*_*h*_(*m*))*_m_* of the non-singleton derived alleles present on that haplotype. In the same way, a separate PCA was performed on each of the 5 continental groups to reveal finescale components of mutation spectrum variation.

### Dating of the TCC*→*T mutation pulse

We estimated the duration and intensity of TCC*→*T rate acceleration in Europe by fitting a simple piecewise-constant rate model to the UK10K frequency data. To specify the parameters of the model, we divide time into discrete log-spaced intervals bounded by time points *t*_1_*, …, t_d_*, assigning each interval a TCC*→*T mutation rate *r*_0_*,…r_d_*. In units of generations before the present, the time discretization points were chosen to be: 20, 40, 200, 400, 800, 1200, 1600, 2000, 2400, 2800, 3200, 3600, 4000, 8000, 12000, 16000, 20000, 24000, 28000, 32000, 36000, 40000. We assume that the total rate *r* of mutations other than TCC*→*T stays constant over time (a first-order approximation).

In terms of these rate variables, we can calculate the expected shape of the TCC*→*T pulse shown in Figure 2B of the main text. The shape of this curve depends on both the mutation rate parameters *r*_*i*_ and the demographic history of the European population, which determines the joint distribution of allele frequency and allele age. To account for the effects of demography, we use Hudson’s ms program to simulate 10,000 random coalescent trees under a realistic European demographic history inferred from allele frequency data [36] and condition our inference upon this collection of trees as follows:

Let *A*(*m, t*) be the function for which 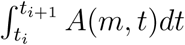 equals the coalescent tree branch length, averaged over the sample of simulated trees, that is ancestral to exactly *m* lineages and falls between time *t*_*i*_ and *t*_*i*+1_. Given this function, which can be empirically estimated from a sample of simulated trees, the expected frequency spectrum entry *k/n* is

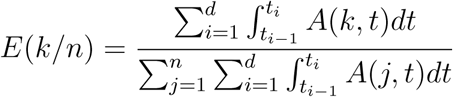

and the expected fraction of TCC*→*T mutations in allele frequency bin *k/n* is

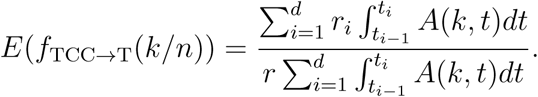

The expected value of the TCC*→*T enrichment ratio being plotted in Figure 2B is

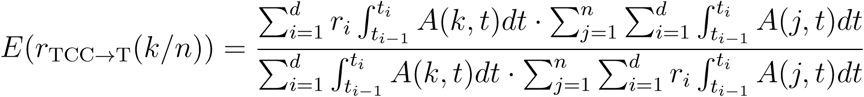

In Figure 2B, enrichment ratios are not computed for every allele frequency in isolation, but for allele frequency bins that each contain similar numbers of SNPs. Given integers 1 *≤ k_m_ < k_m_*+1 *≤ n*, the expected TCC*→*T enrichment ratio averaged over all SNPs with allele frequency between *k*_*m*_*/n* and *k*_*m*+1_*/n* is:

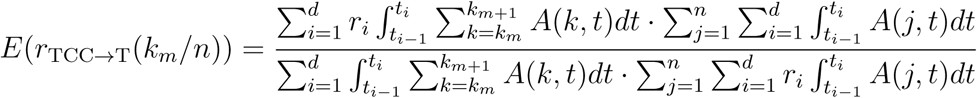

We optimize the mutation rates *r*_1_*, …, r_d_* using a log-spaced quantization of allele frequencies *k*_1_*/n, …, k_m_/n* defined such that all bins contain similar numbers of SNPs. The chosen allele count endpoints *k*_1_*, …, k_m_* are: 1, 2, 3, 4, 5, 6, 7, 8, 9, 10, 20, 30, 40, 50, 60, 70, 80, 90, 100, 200, 300, 400, 500, 600, 700, 800, 900, 1000, 2000, 3000, 4000. Given this quantization of allele frequencies, we optimize *r*_1_*, …, r_d_* by using the BFGS algorithm to minimize the least squares distance *D*(*r*_0_*, …, r_d_*) between *E*(*r*_TCC_*→*T(*k_m_/n*)) and the empirical ratio *r*_TCC_*→*T(*k_m_/n*) computed from the UK10K data. This optimization is subject to a regularization penalty that minimizes the jumps between adjacent mutation rates *r_i_* and *r*_*i*+1_:

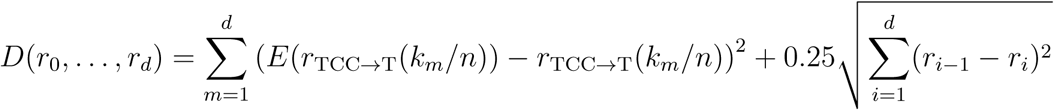

Although the underlying model of mutation rate change assumed here is very simple, it still represents an advance over the method used in [12] to estimate of the timing of the TCC*→*TTC mutation rate increase. That method relied upon explicit estimates of allele age from a dataset of less than 100 individuals, which are much noisier than integration of a joint distribution of allele age and frequency across a sample of thousands of haplotypes.

## Supplementary Figures

**Figure 1– Figure Supplement 1:**
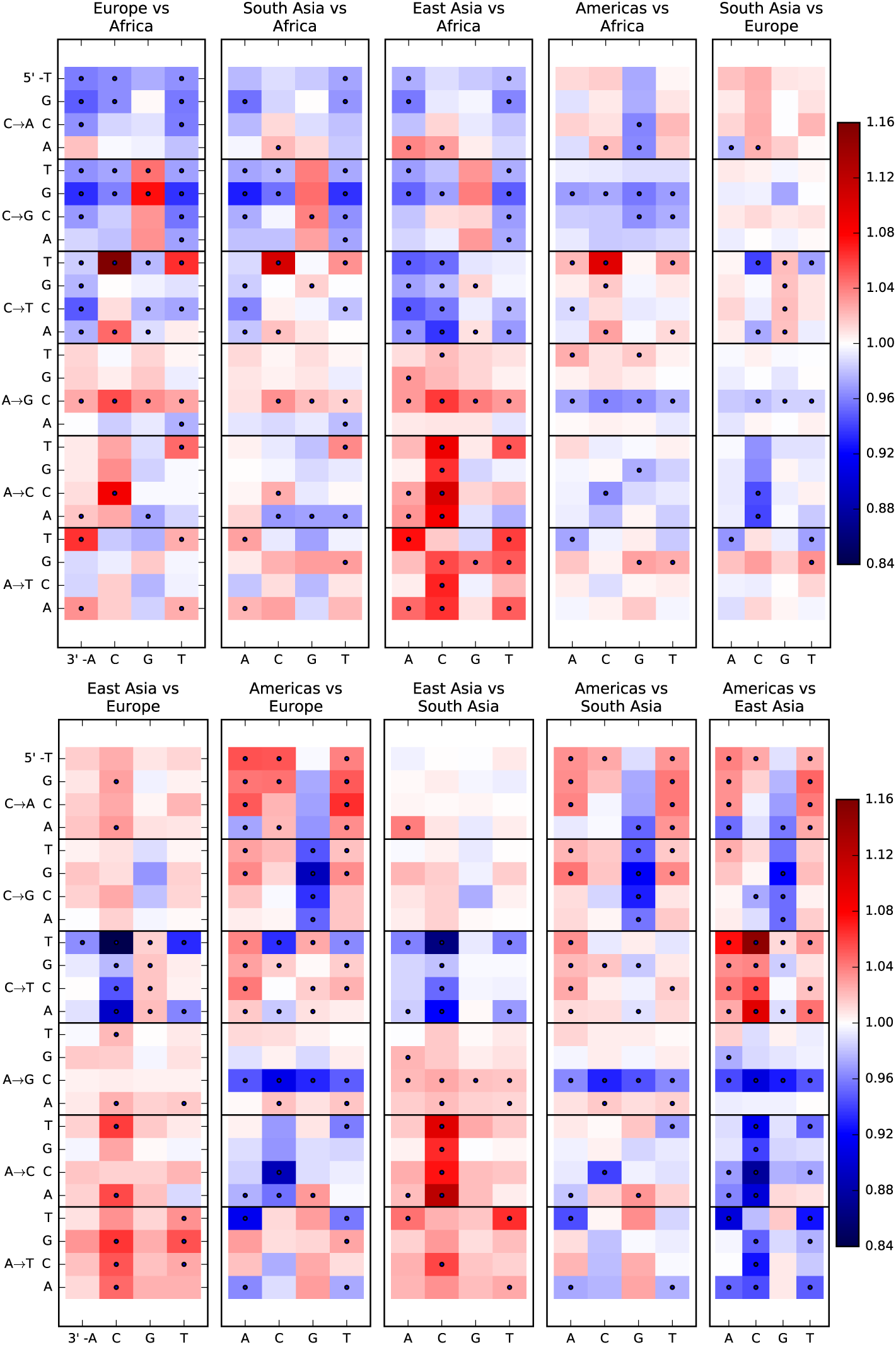
Pairwise mutation spectrum comparisons among continental groups. Each of these plots compares the mutation spectra of two populations *P*_1_ and *P*_2_. Letting *f_i_* denote the fraction of SNVs in population *P_i_* that have a given triplet context, ancestral allele, and derived allele, the corresponding heat map square visualizes the enrichment ratio *f*_1_*/f*_2_. Black dots mark mutation types for which the difference between populations has a ^2^ *p*-value less than 10*−*^5^.

**Figure 1–Figure Supplement 2:**
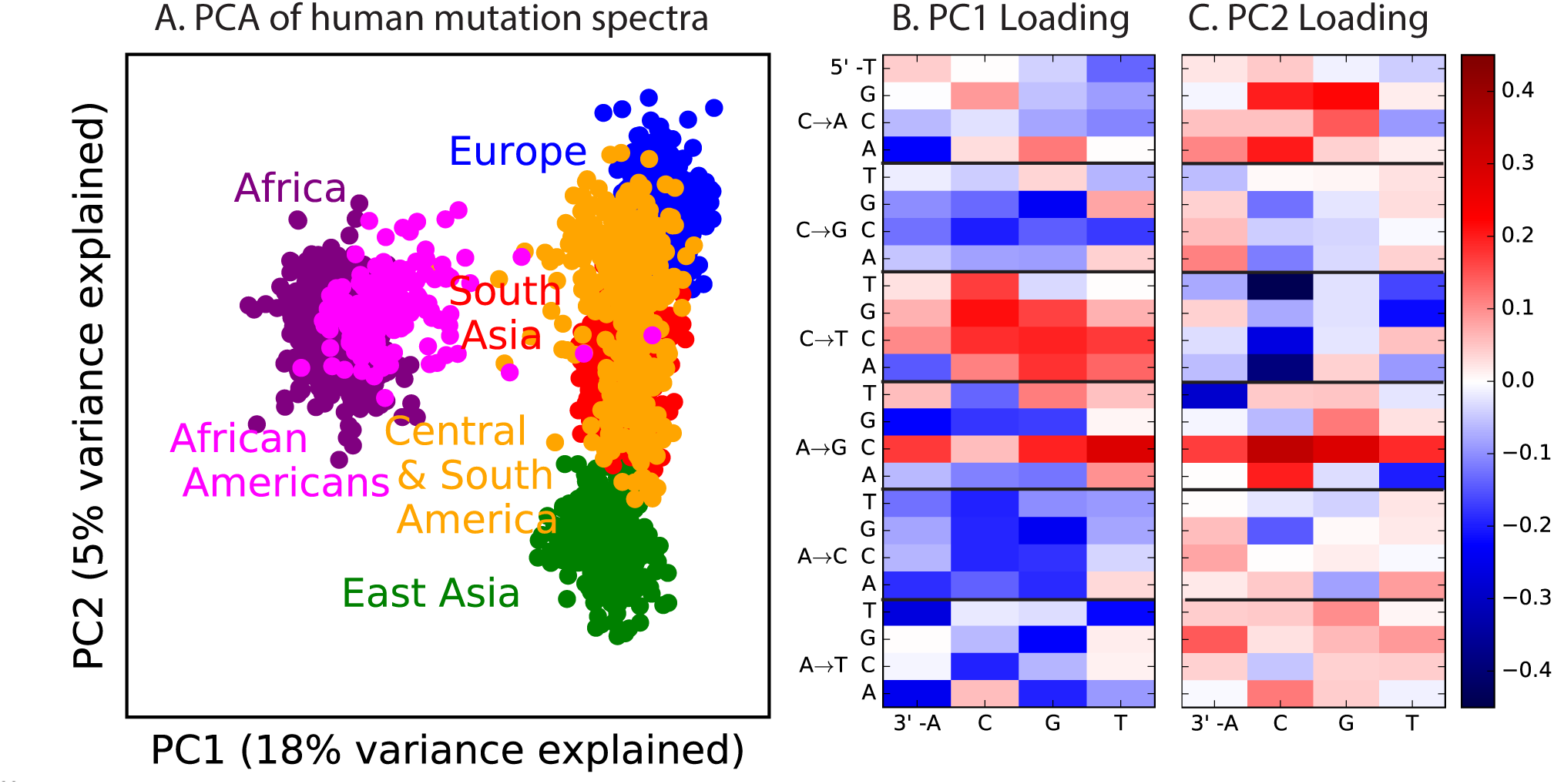
PCA of all 1000 Genomes continental groups. All admixed North and South American individuals were omitted from Figure 1 in the main text to clarify the separation of other populations along an African vs non-African axis and an East vs West Eurasian axis. Here, admixed Americans are added in black. As expected, some African-Americans group with the Africans, while other admixed Americans fall within the variation of other East and West Eurasians. The accompanying heat maps show the mutation type loadings of the first two principal components, the second of which is heavily weighted toward the European TCC*→*TTC signature.

**Figure 1–Figure Supplement 3:**
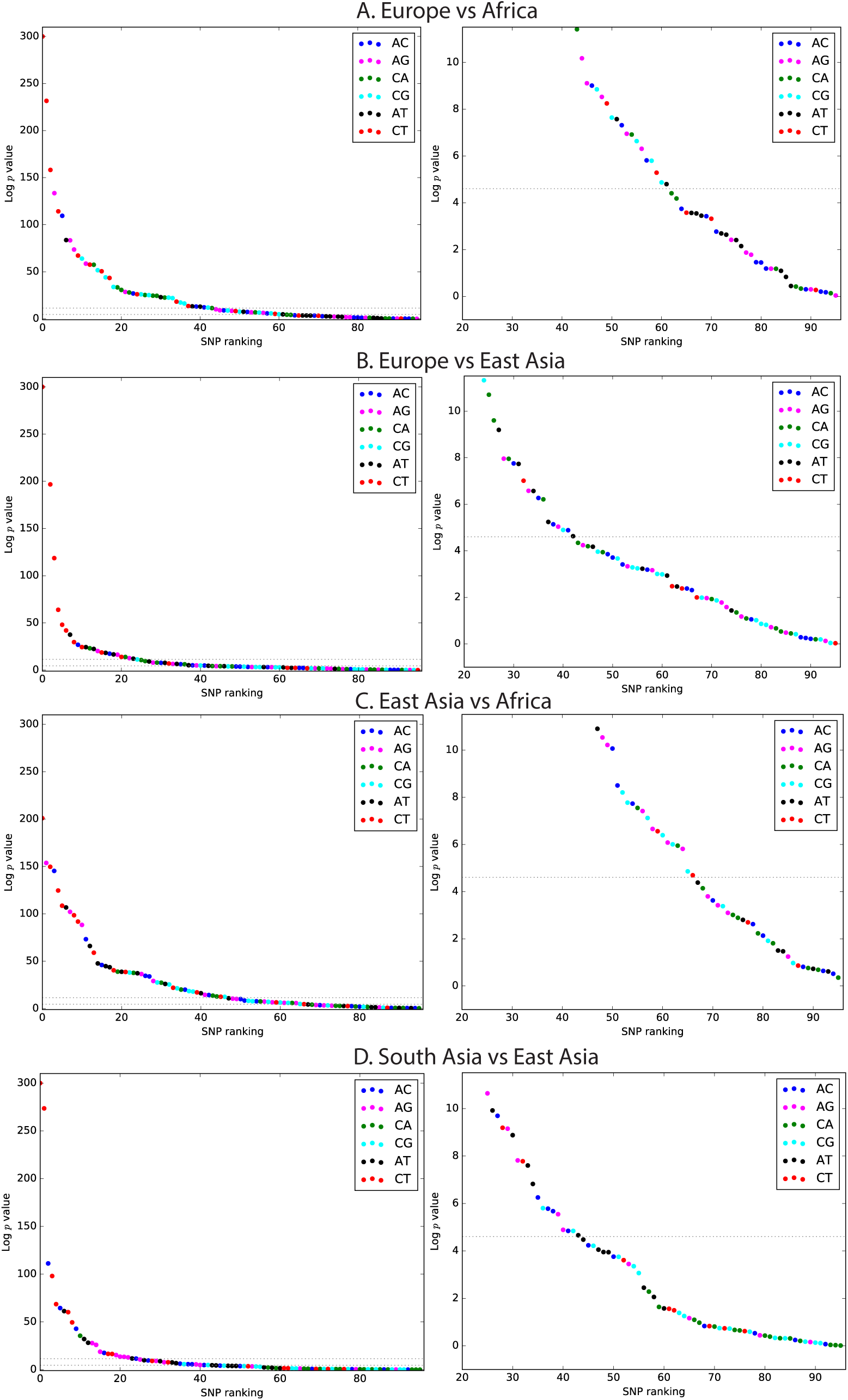
Mutation spectrum comparison *p*-values. Each left-hand plot shows all chi-squared *p*-values corresponding to the ratios from Figure 1A. In the absence of recent mutation spectrum evolution, only one out of 96 SNP categories is expected to have a *p*-value below 0.01 (lower dotted line). In contrast, the majority of *p* values meet the more stringent threshold *p <* 1*e −* 5. The corresponding right hand panel shows a closeup of the distribution of *p*-values greater than 1*e −* 5.

**Figure 1–Figure Supplement 4:**
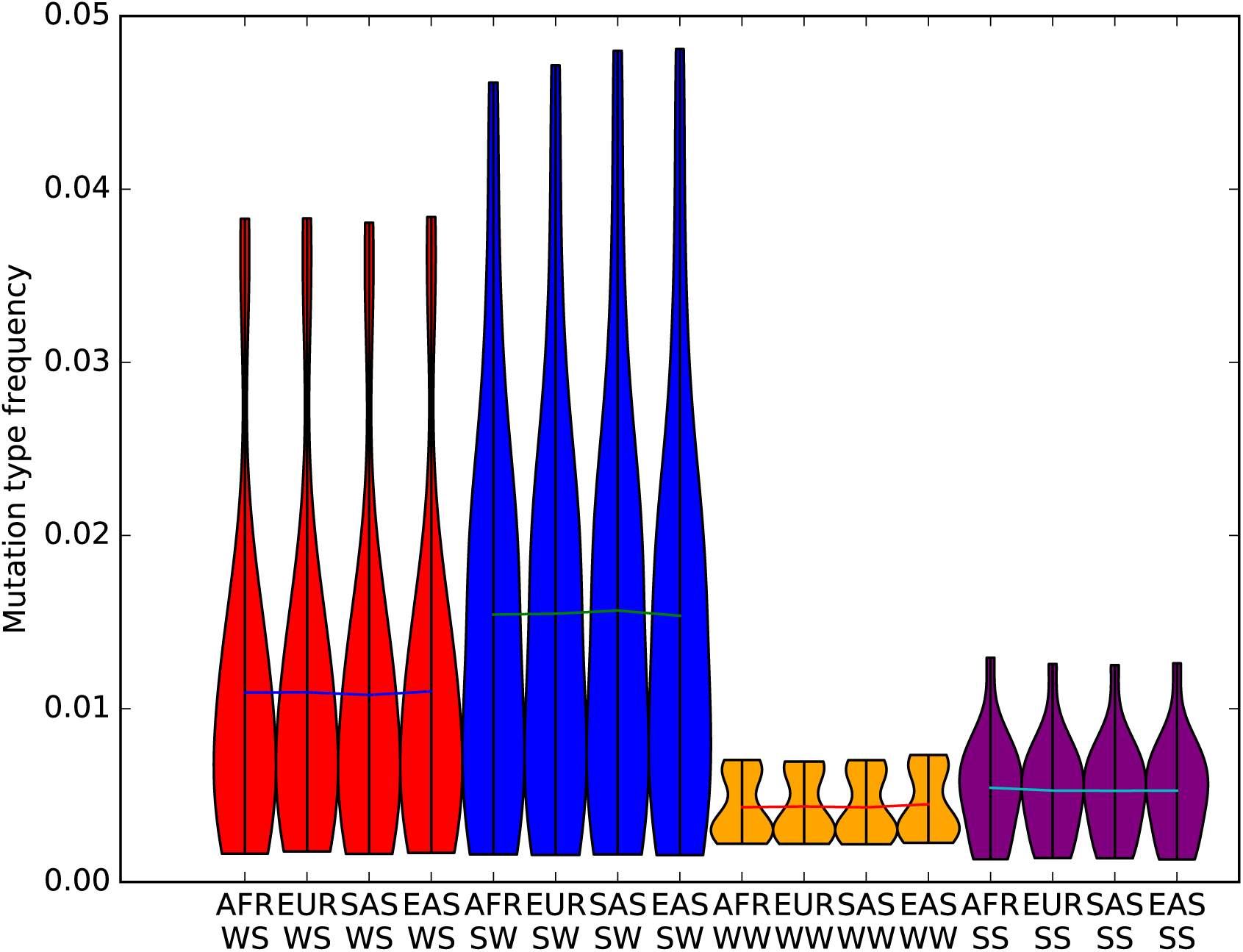
The effects of biased gene conversion on mutation spectra. When using segregating variation to study the mutation spectrum, one potential source of bias is that strong-to-weak mutations, where the ancestral allele is G or C and the derived allele is A or T, have a lower fixation probability than weak-to-strong mutations due to biased gene conversion (BGC). If this effect were sufficiently strong, it would inflate the apparent mutation fractions of weak-to-strong mutations, especially in populations with large effective sizes where natural selection is particularly efficient. Within humans, Africans have the largest long-term effective population size, while East Asians and Native Americans have the lowest. Therefore, if BGC has created differences in mutation spectra between populations, the fraction of weak-to-strong SNVs should be highest in Africans, intermediate in Europeans and South Asians, and lowest in East Asians and Native Americans. This violin plot reveals no such pattern, suggesting that BGC is not a strong driver of mutation spectrum differences between human populations. We do not observe either a direct correlation between in strong-to-weak mutation fraction and distance from Africa or an inverse correlation between weak-to-strong mutation fraction and distance from Africa.

**Figure 1–Figure Supplement 5:**
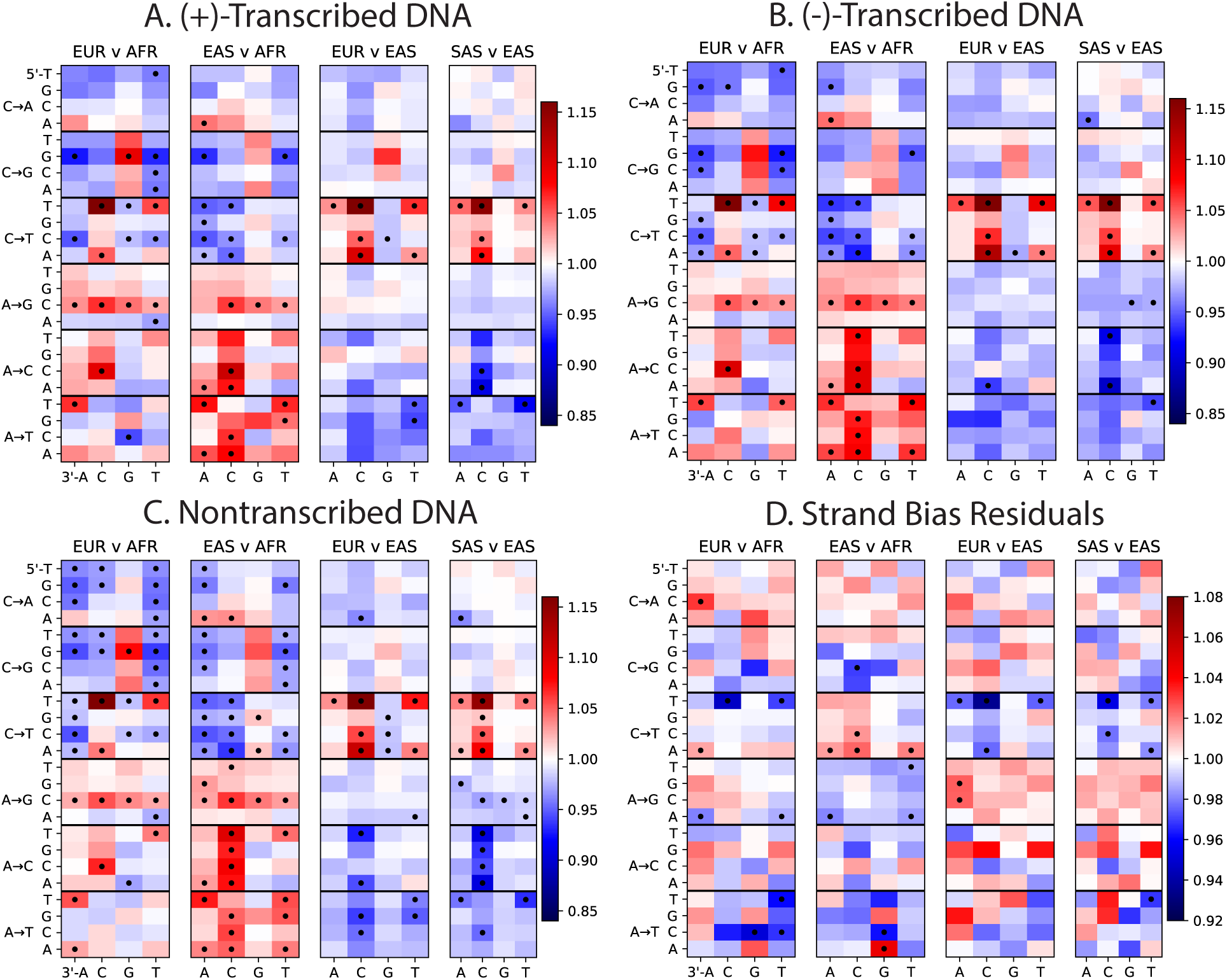
Mutation Spectra of Transcribed vs Non-Transcribed DNA. Using the UCSC Genome Browser annotations of the human reference hg19, we determined whether each SNP occurs in a transcribed or non transcribed region. We further divided SNPs occurring in transcribed regions according to whether the ancestral A or C allele occurs on the (+)-strand or the (-)-strand. Panels A, B, and C all show the same population-specific mutation type enrichments that are observed in Figure 1B. Panel D plots the residuals between panels A and B, highlighting mutation types that show a modest difference in strand bias between populations.

**Figure 1–Figure Supplement 6:**
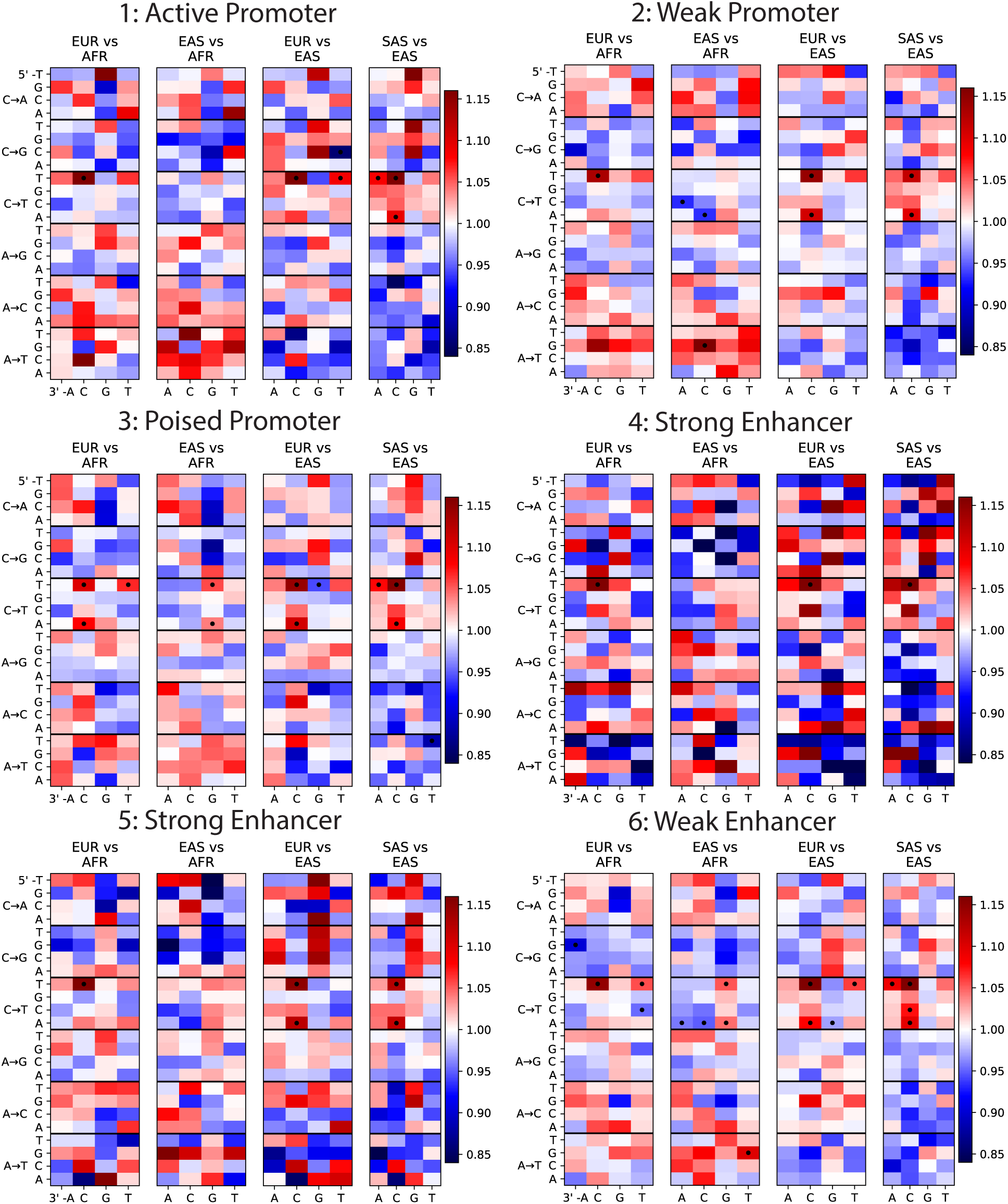
Mutation Spectra of ChromHMM chromatin states (Part I of II). To investigate whether any mutation spectrum shifts might be confined to particular chromatin states, we used chromHMM annotations of the human embryonic stem cell line HESC-H1 [37]. Each heat map plots mutation spectrum comparisons for SNPs that are annotated as being part of the same chromatin state, and dots mark mutation types that show a significant enrichment in one population at the level *p <* 0.01. Every chromatin state shows enrichment of the TCC*→*TTC signature in Europe and South Asia. Some heat maps are noisy due to the small sample size of SNPs contained within these regions, but all showcase the same general patterns as Figure 1B.

**Figure 1–Figure Supplement 7:**
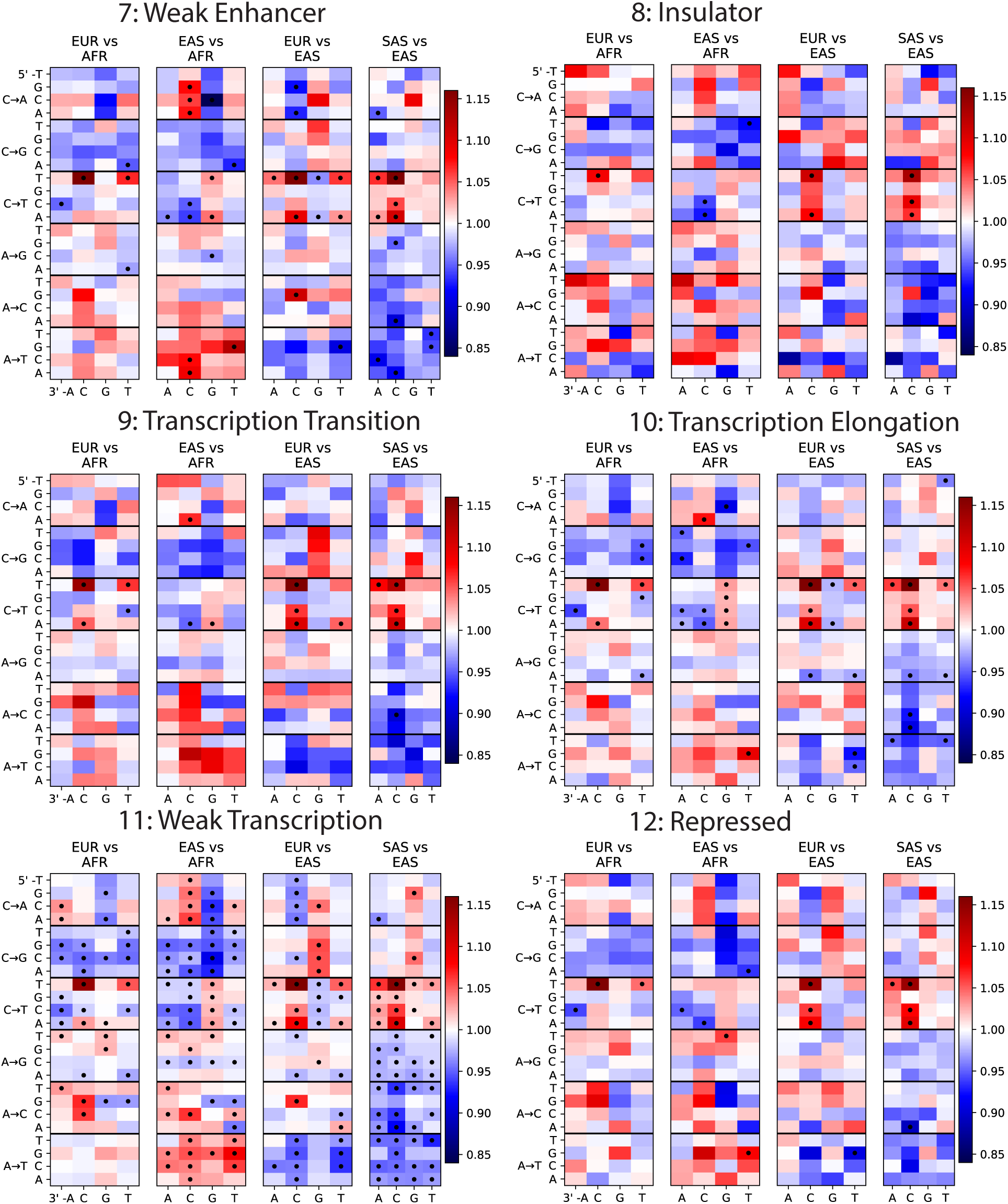
Mutation Spectra of ChromHMM chromatin states (Part II of II).

**Figure 1–Figure Supplement 8:**
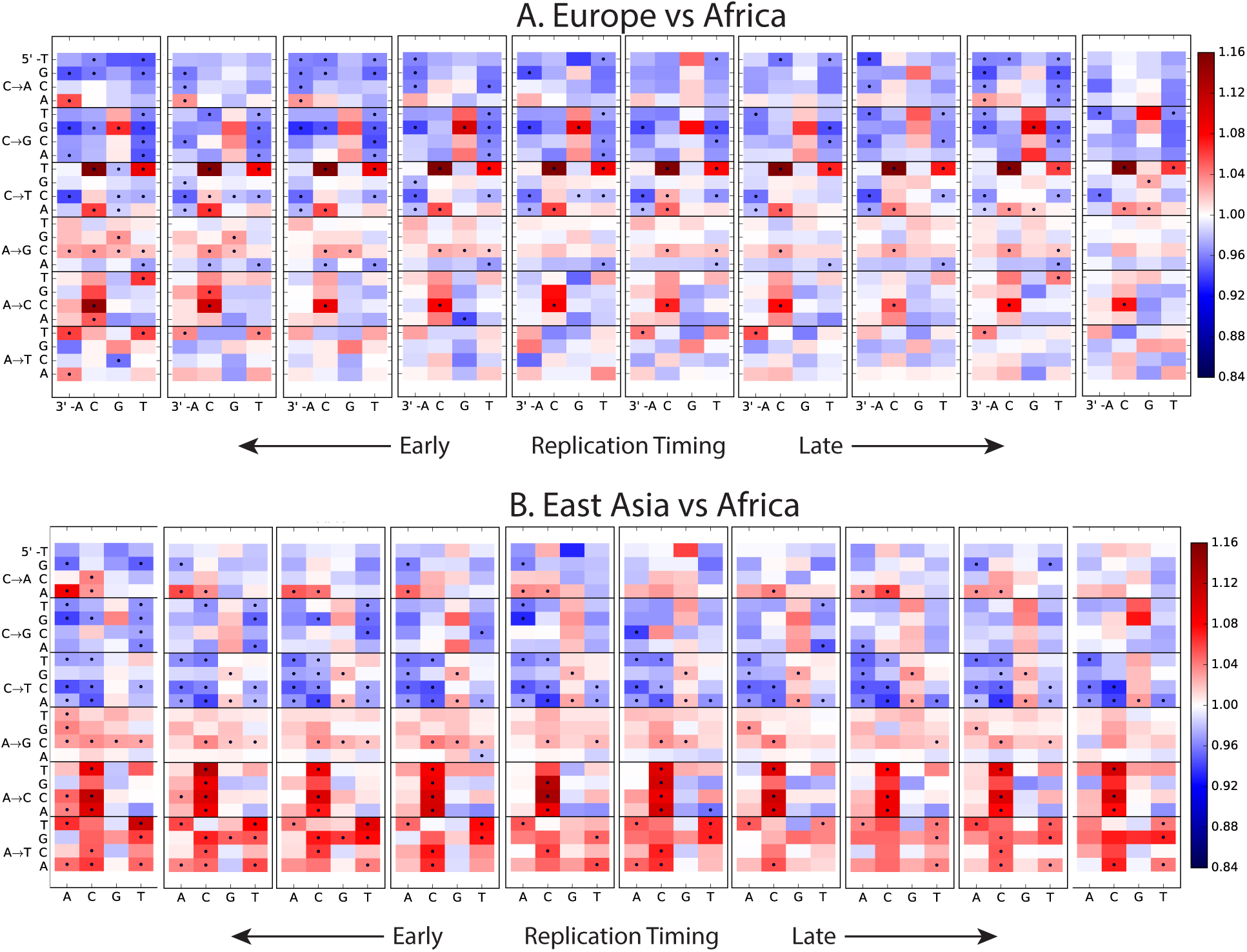
Variation of the mutation spectrum with DNA replication timing. We partitioned the genome into 10 equal replication timing quantiles using data obtained from [38], then computed mutation spectrum differences within each quantile. Although most patterns from Figure 1B replicate within each replication timing bin, there are a few exceptions. CpG transitions, which occur most often in early-replicating regions, vary in population bias depending on replication timing. In addition, the deficit of ACA*→*AAA and AAA*→*ATA mutations in Africa compared to Europe and Asia is observed mainly in early-replicating regions.

**Figure 1–Source Data 1.** This text file shows the number of SNPs in each of the 96 mutational categories that passed all filters in each 1000 Genomes continental group.

**Figure 2–Figure Supplement 1:**
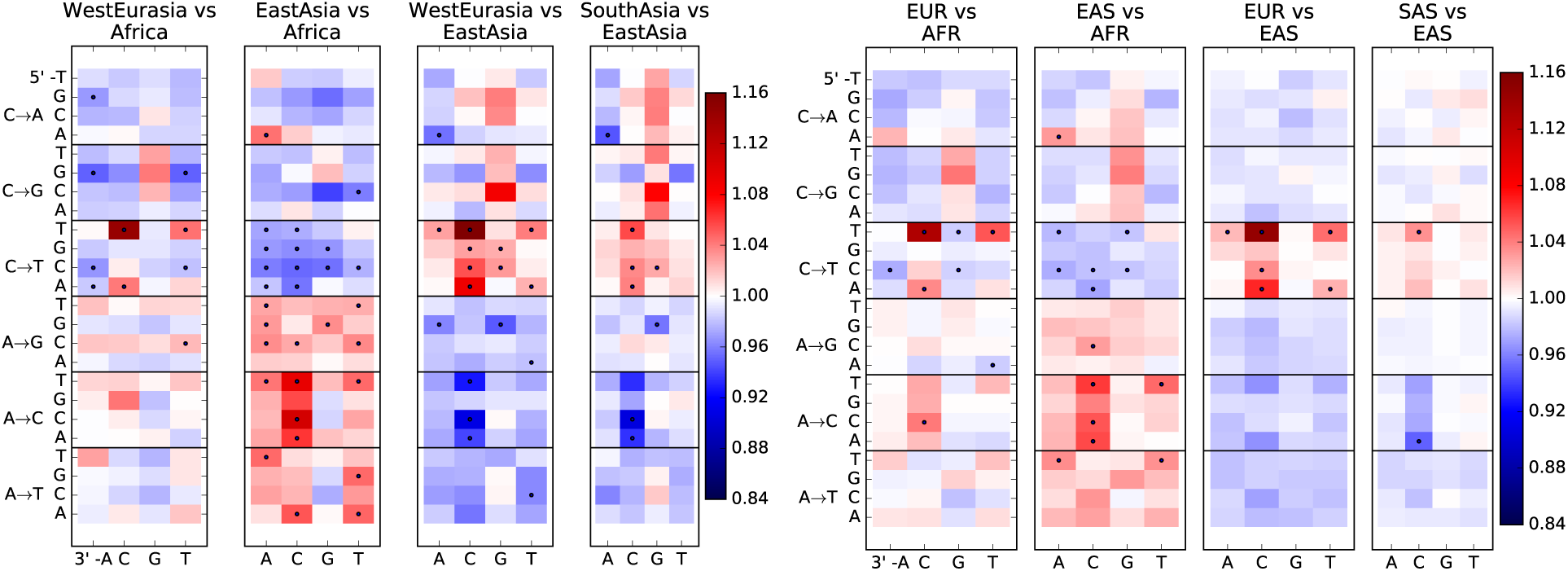
Heatmap comparisons between continental groups in 1000 Genomes and the SGDP. Here, each 1000 Genomes population is projected down to the sample size of the corresponding SGDP population in order to sample alleles with a similar distribution of ages and frequencies.

**Figure 2–Figure Supplement 2:**
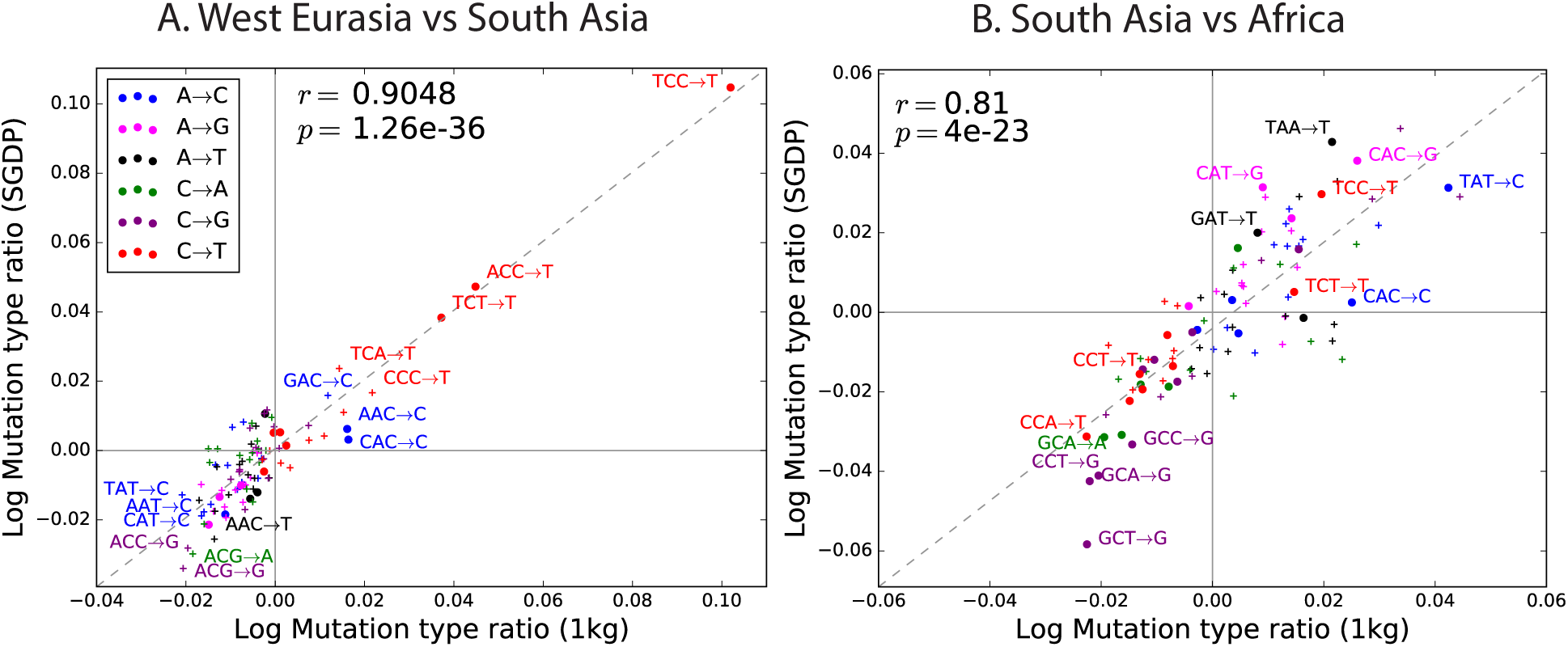
Regression of the SGDP heatmap coefficients versus the corresponding 1000 Genomes heatmap coefficients.

**Figure 3–Figure Supplement 1:**
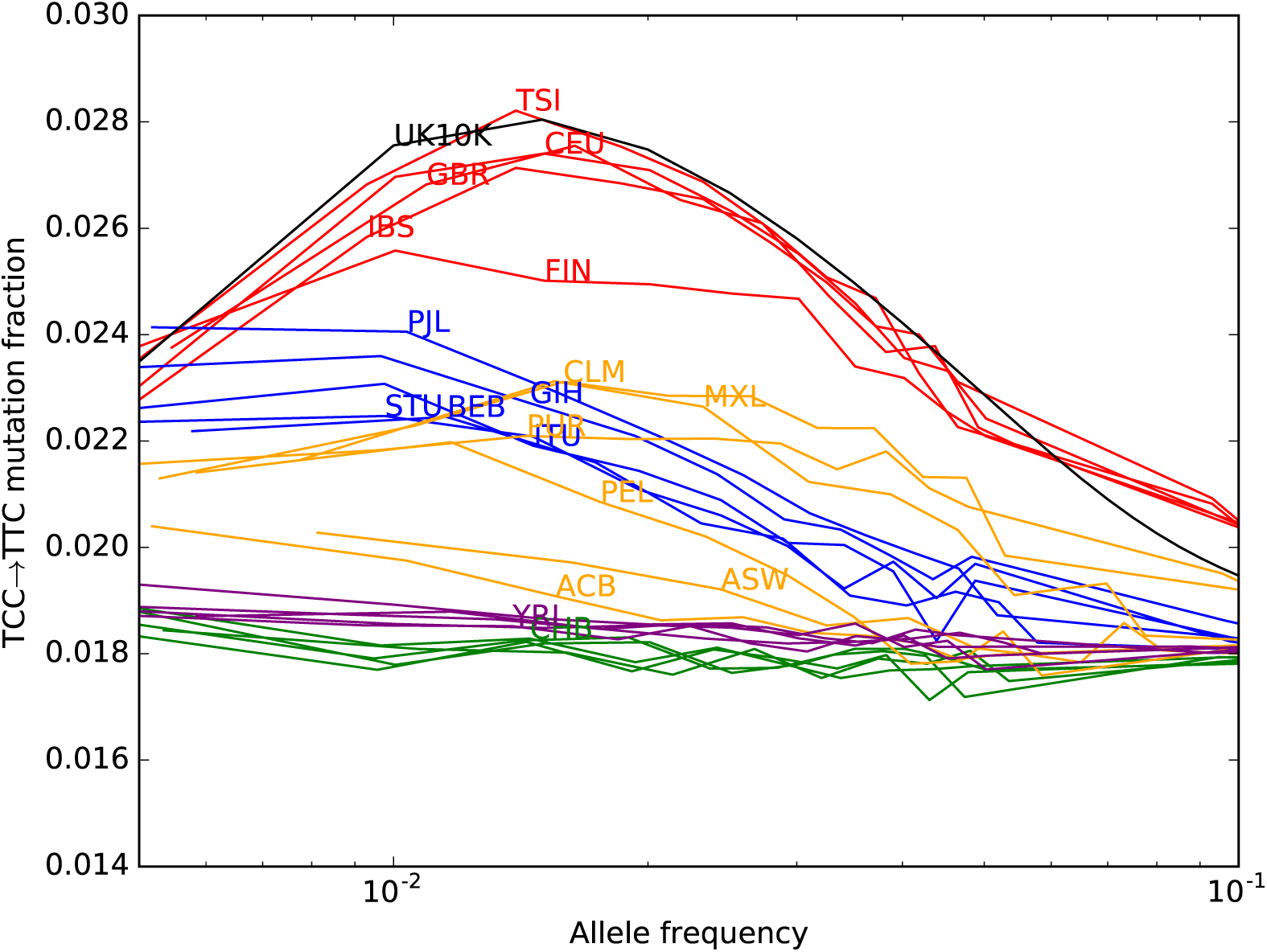
TCC→TTC mutation fraction as a function of allele frequency in all 1000 Genomes populations. To enable better comparison with the 1000 Genomes data, the UK10K SNPs have been downsampled to 200 individuals. The age distribution of alleles of a given frequency varies as a function of the number of lineages being sampled–this is why the UK10K pulse peaks around 0.6% frequency when measured in a dataset of thousands of lineages, but peaks around 2% in a subsample of only 400 lineages. Some African and East Asian population names have been omitted for clarity since the TCC*→*TTC mutation fraction is so uniform within these continental groups. Red= European populations; Blue = South Asian; Orange = Americas; Purple = Africa; Green = East Asia.

**Figure 3–Figure Supplement 2:**
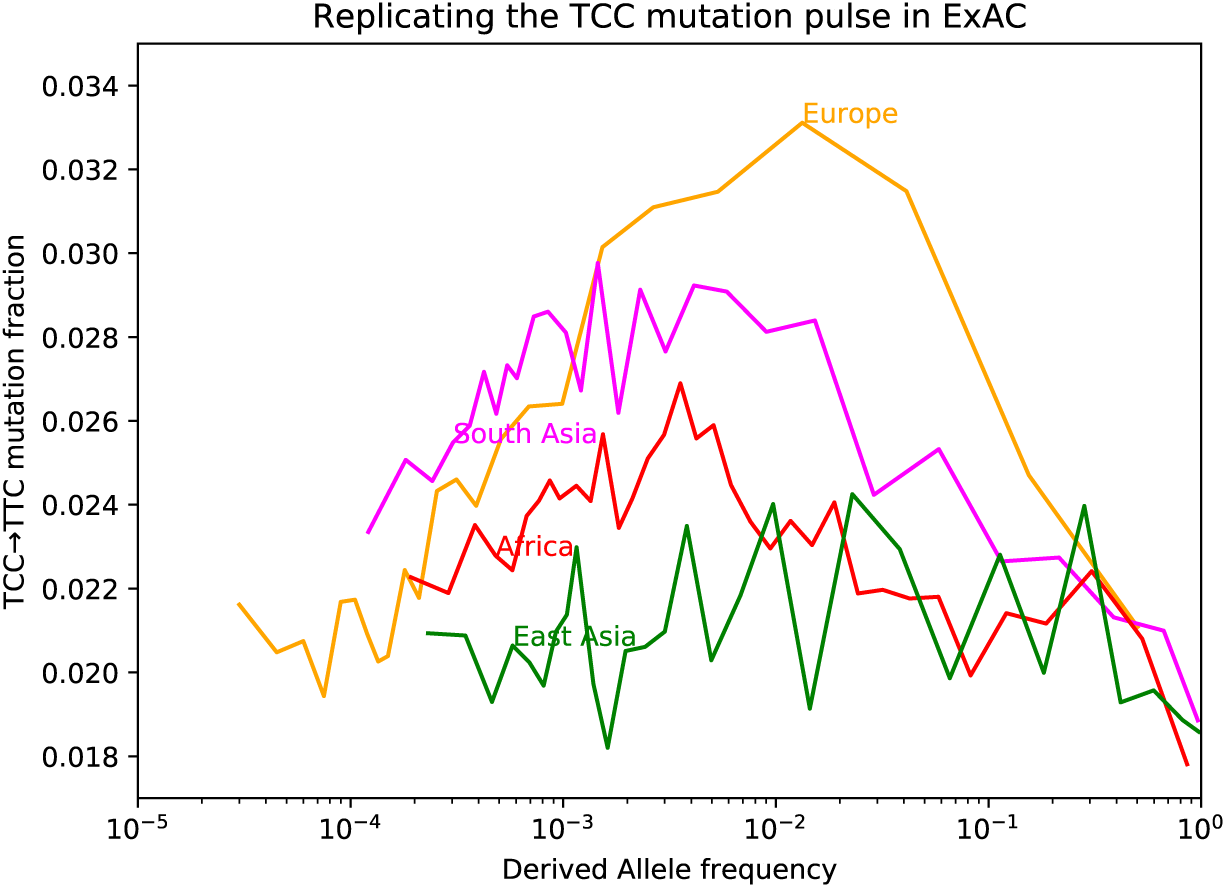
Fraction of TCC*→*TTC mutations as a function of allele frequency in ExAC. Lek, et al. compiled data from 60,706 exomes to create the Exome Aggregation Consortium dataset, which enables the analysis of ultra-rare human variation [39]. The overall fraction of TCC*→*TTC mutations is slightly higher in exome data than in whole genome data because exons contain a skewed distribution of triplet contexts, but the pulse pattern from Figure 3B reproduces unmistakably.

**Figure 3–Figure Supplement 3:**
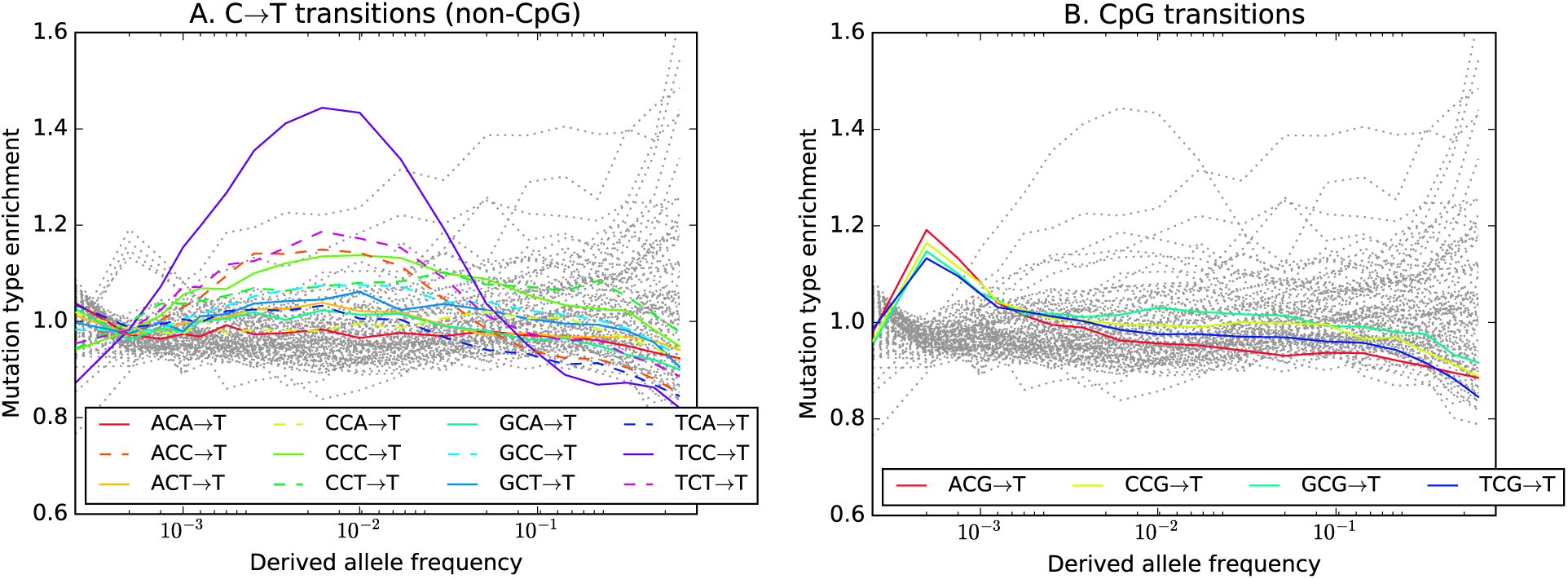
Mutation type enrichment as a function of allele frequency in UK10K (Part I of III). The eleven panels in Figure Supplements 2, 3, and 4 show the full dependence of mutation spectrum on allele frequency in the UK10K data. If we let *F* (*f, m*) denote the fraction of SNVs of frequency *f* that are of type *m* and let *F* (*m*) denote the fraction of all mutations that are of type *m*, the enrichment of mutation type *m* as a function of frequency is *F* (*f, m*)*/F* (*m*). This function is expected to fluctuate around *y* = 1 unless the rate of *m* has recently increased or decreased. All 96 mutation types are visualized in every panel, but most corresponding lines are greyed out to enhance readability. Some lines deviate from *y* = 1 due to the effects of biased gene conversion (BGC)–this occurs when one of the ancestral or derived alleles is a weak base (A or T, abbreviated W) and the other allele is a strong base (G or C, abbreviated S). W*→*S mutations are more abundant at high allele frequencies, while S*→*W mutations are more abundant at low frequencies. These effects are visible but modest in panels D, G, H, and I, but much more pronounced in panels B, C, and F, which focus on mutations in the CpG context. Transitions of the type CpA*→*CpG, which create CpG motifs, are extremely enriched at high frequencies, and this pattern may be an artifact of ancestral misidentification [40]. CpG motifs have such high mutation rates that CpG*→*CpT transitions often happen at the same site in humans and chimps, and these low-frequency double mutations are misclassified as high-frequency CpT*→*CpG mutations. Although it is not surprising to see a peak of CpT*→*CpG transitions at high frequencies in panel F, it is somewhat surprising to see CpG*→*GpG transversions peak in abundance at high frequencies in panel C. This might be a signature of recent declines in the rates of these mutations, since neither ancestral misidentification nor biased gene conversion is thought to produce such a pattern. In addition, neither of these processes can explain the strong enrichment of certain A*→*T mutations at high frequencies that is observed in panel K.

**Figure 3–Figure Supplement 4:**
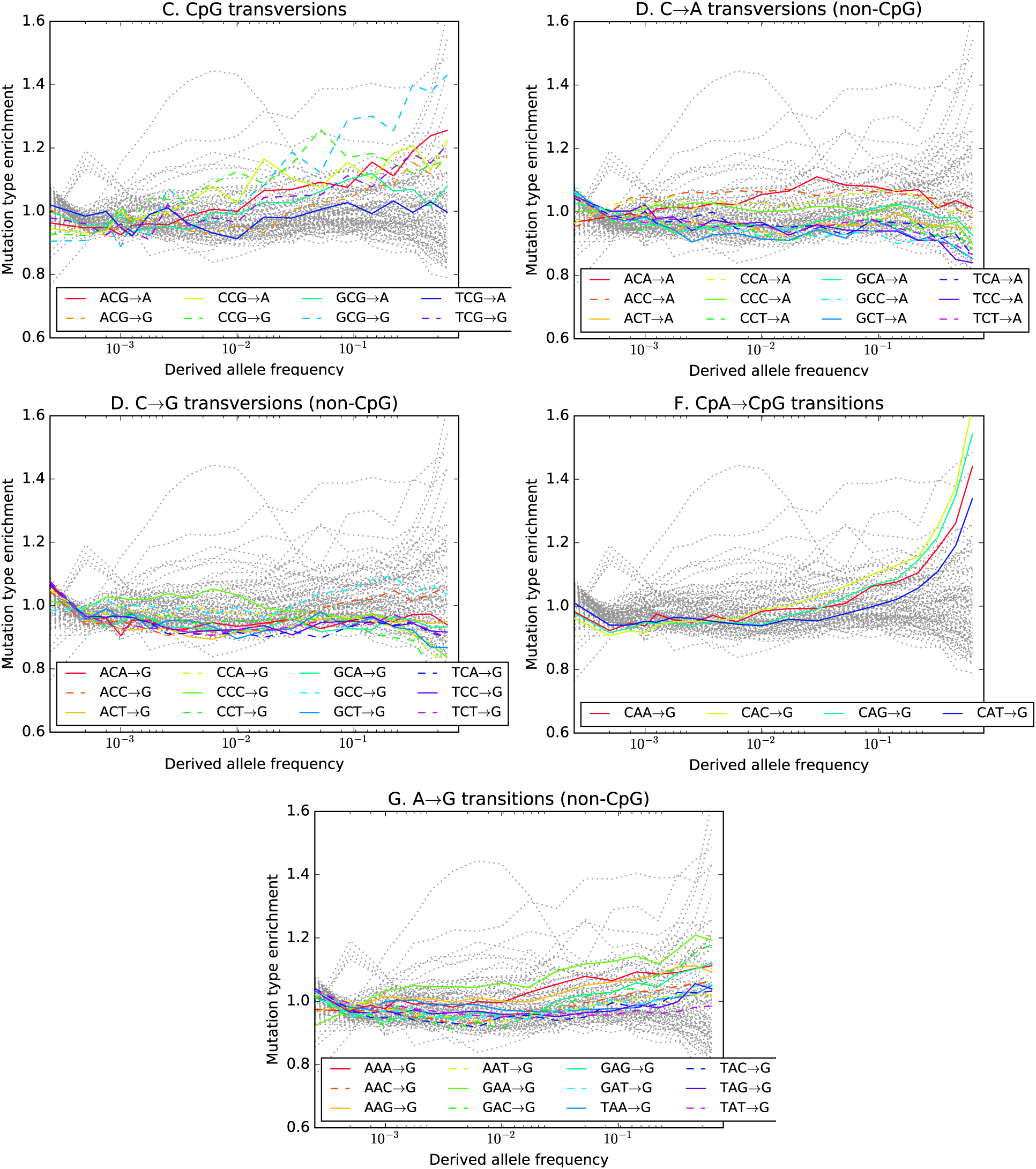
Mutation type enrichment as a function of allele frequency in UK10K (Part II of III). The eleven panels in this 3-part figure show the full dependence of mutation spectrum on allele frequency in the UK10K data.

**Figure 3–Figure Supplement 5:**
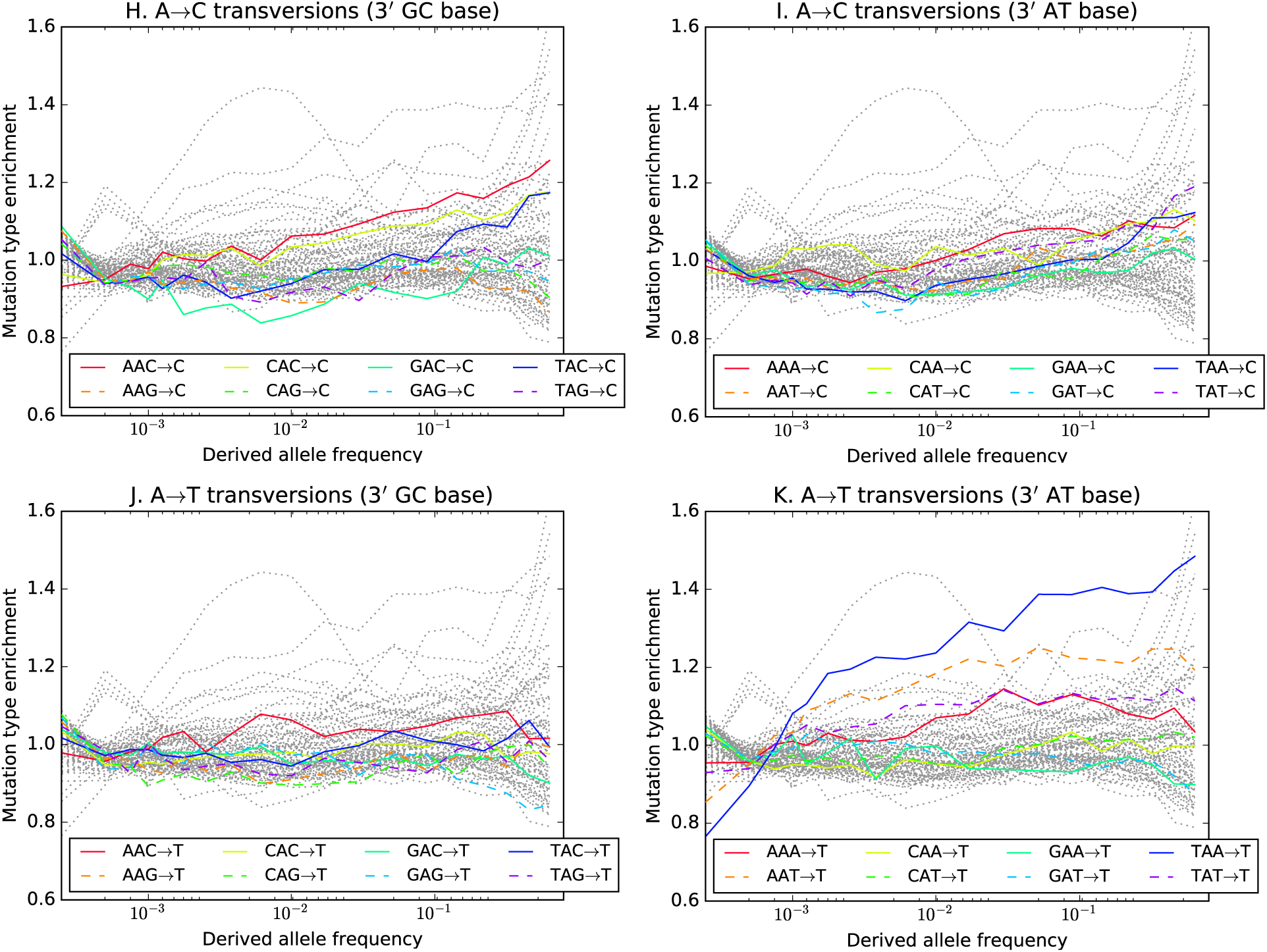
Mutation type enrichment as a function of allele frequency in UK10K (Part III of III). The eleven panels in this 3-part figure show the full dependence of mutation spectrum on allele frequency in the UK10K data.

**Figure 3–Figure Supplement 6:**
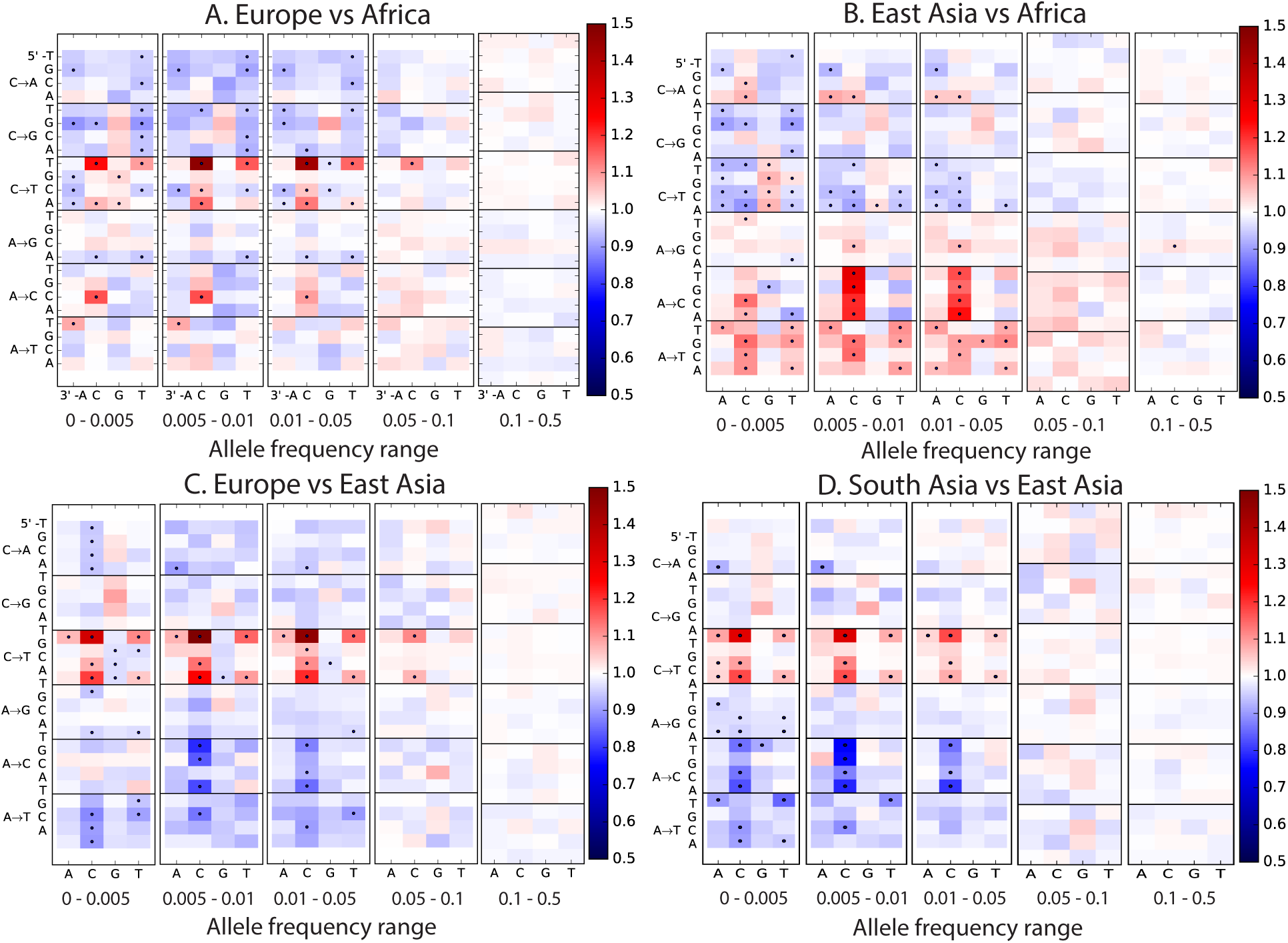
Mutation spectrum comparisons partitioned by allele frequency. Each of these heatmaps shows a subset of the data used to construct Figure 1B, partitioned by allele frequency to show how rare variants are the most highly differentiated between populations. Black dots highlight mutation types that are significantly different in abundance between two populations in a particular frequency class at the *p <* 10*−*^5^ level according to a chi-square test.

**Figure 4–Figure Supplement 1:**
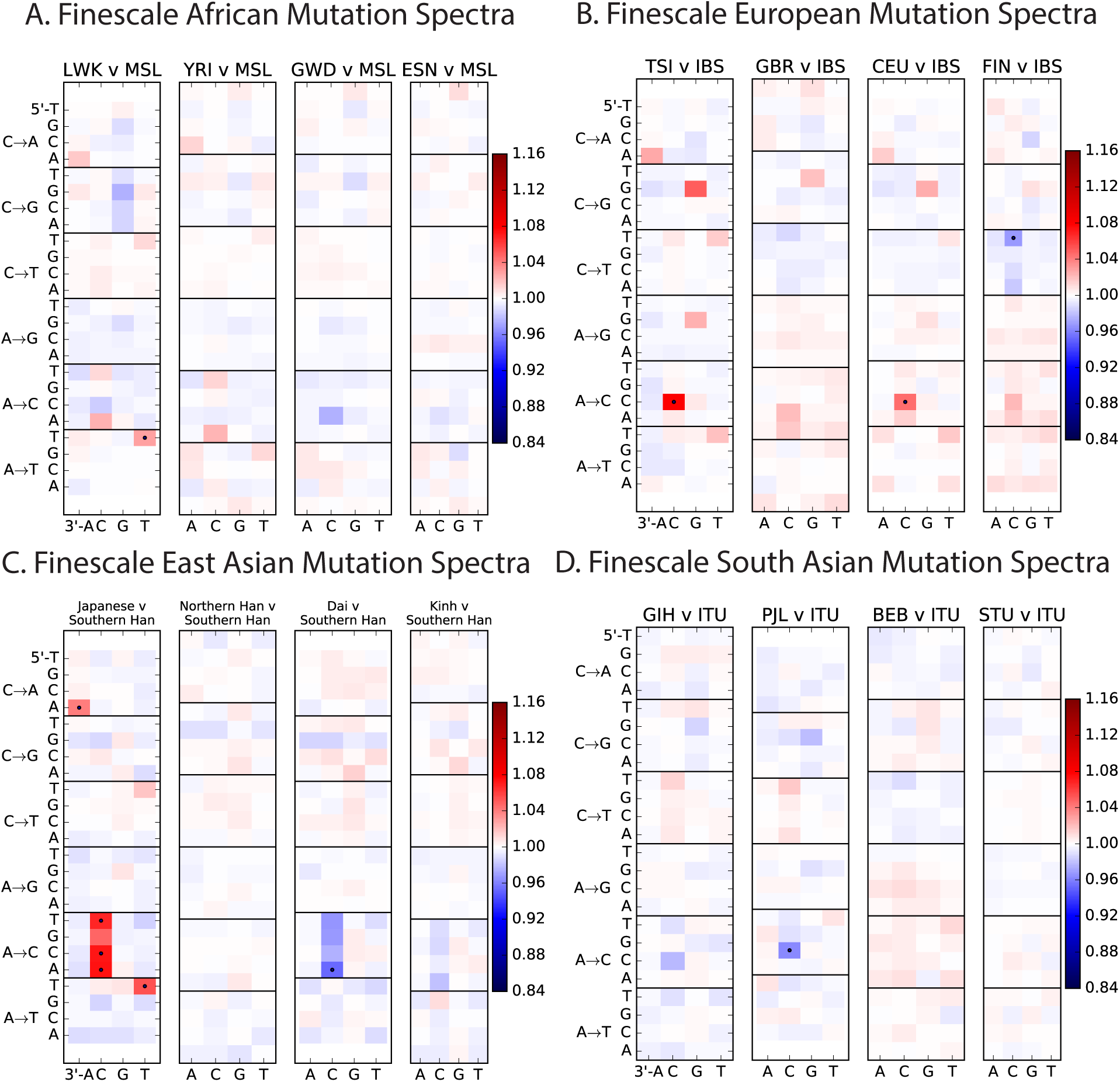
Mutation spectrum differences within Africa, Europe, East Asia, and South Asia. Figure 4B of the main text shows heat map comparisons between East Asian populations, which display fine-scale differences that are exceptionally well defined. For completeness, this figure shows finescale heatmap comparisons within all 1kG continental groups. We can see that CAC*→*CCC and TAT*→*TTT are heterogeneously distributed within multiple continents, but to the greatest extent in East Asia. In addition the TCC*→*TTC signature is somewhat heterogeneously distributed within Europe and South Asia, being depleted in Finns and enriched in the Punjabi and Gujarati. Each continental group in the 1000 Genomes data is divided into 5 sub-populations. These heat maps compare the mutation spectra of these fine-scale populations to each other. African populations are: MSL = Mende in Sierra Leone; LWK = Luhya in Webuye, Kenya; YRI = Yoruba in Ibadan, Nigeria; GWD = Gambian in Western Divisions; ESN = Esan in Nigeria. European populations are: IBS = Iberian Population in Spain; TSI = Toscani in Italia; GBR = British in England and Scotland; CEU = Utah Residents (CEPH) with Northern and Western Ancestry; FIN = Finnish in Finland. East Asian populations are: CDX = Chinese Dai in Xishuangbanna, China; JPT = Japanese in Tokyo, Japan; CHB = Han Chinese in Bejing, China; CHS = Southern Han Chinese; KHV = Kinh in Ho Chi Minh City, Vietnam. South Asian populations are: ITU = Indian Telugu from the UK; GIH = Gujarati Indian from Houston, Texas; PJL = Punjabi from Lahore, Pakistan; BEB = Bengali from Bangladesh; STU = Sri Lankan Tamil from the UK.

**Figure 4–Figure Supplement 2:**
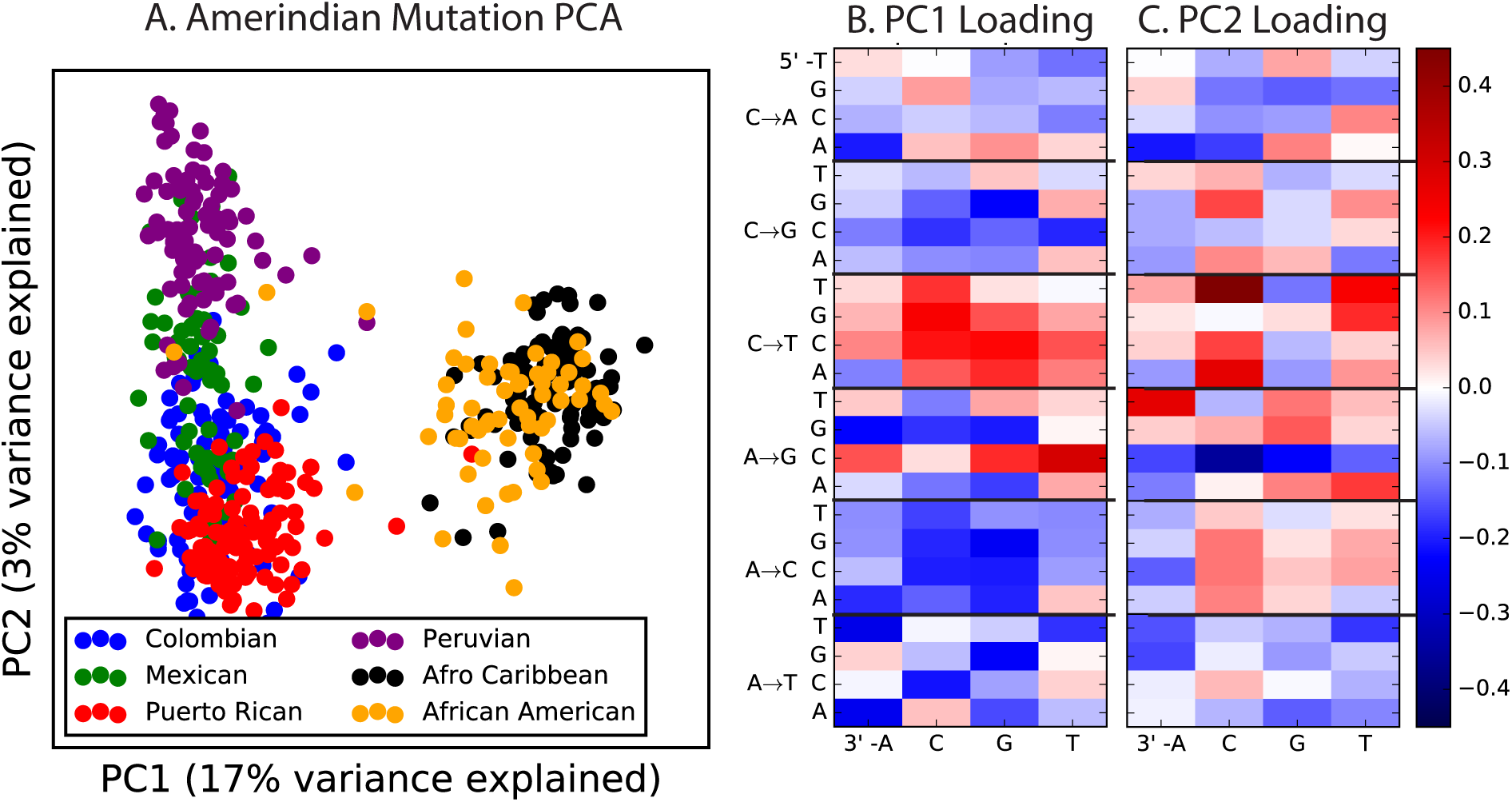
PCA of American populations. Population abbreviations are: CLM = Colombians from Medellin, Colombia; MXL = Mexican Ancestry from Los Angeles, USA; PUR = Puerto Ricans from Puerto Rico; PEL = Peruvians from Lima, Peru; ACB = African Caribbeans in Barbados; ASW = Americans of African Ancestry in SW USA. Admixed populations from the Americans show structure that mirrors the continental groups, with PC1 essentially measuring the ratio between African and non-African ancestry and PC2 measuring the ratio between European and Native American ancestry. The accompanying heat maps show the loadings of the first two principal components.

**Figure 4–Figure Supplement 3:**
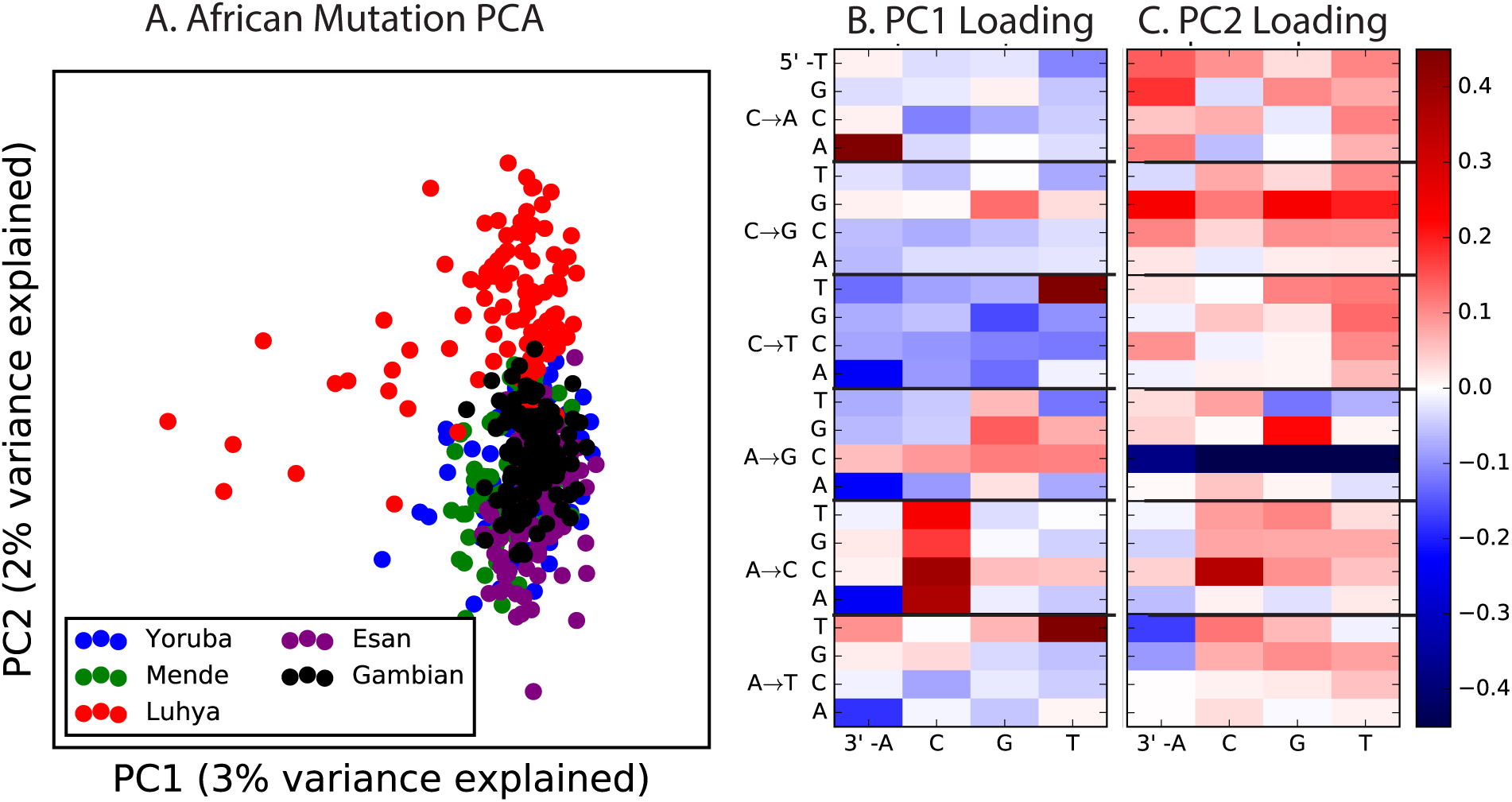
PCA of African populations. Population abbreviations are: MSL = Mende in Sierra Leone; LWK = Luhya in Webuye, Kenya; YRI = Yoruba in Ibadan, Nigeria; GWD = Gambian in Western Divisions; ESN = Esan in Nigeria.

**Figure 4–Figure Supplement 4:**
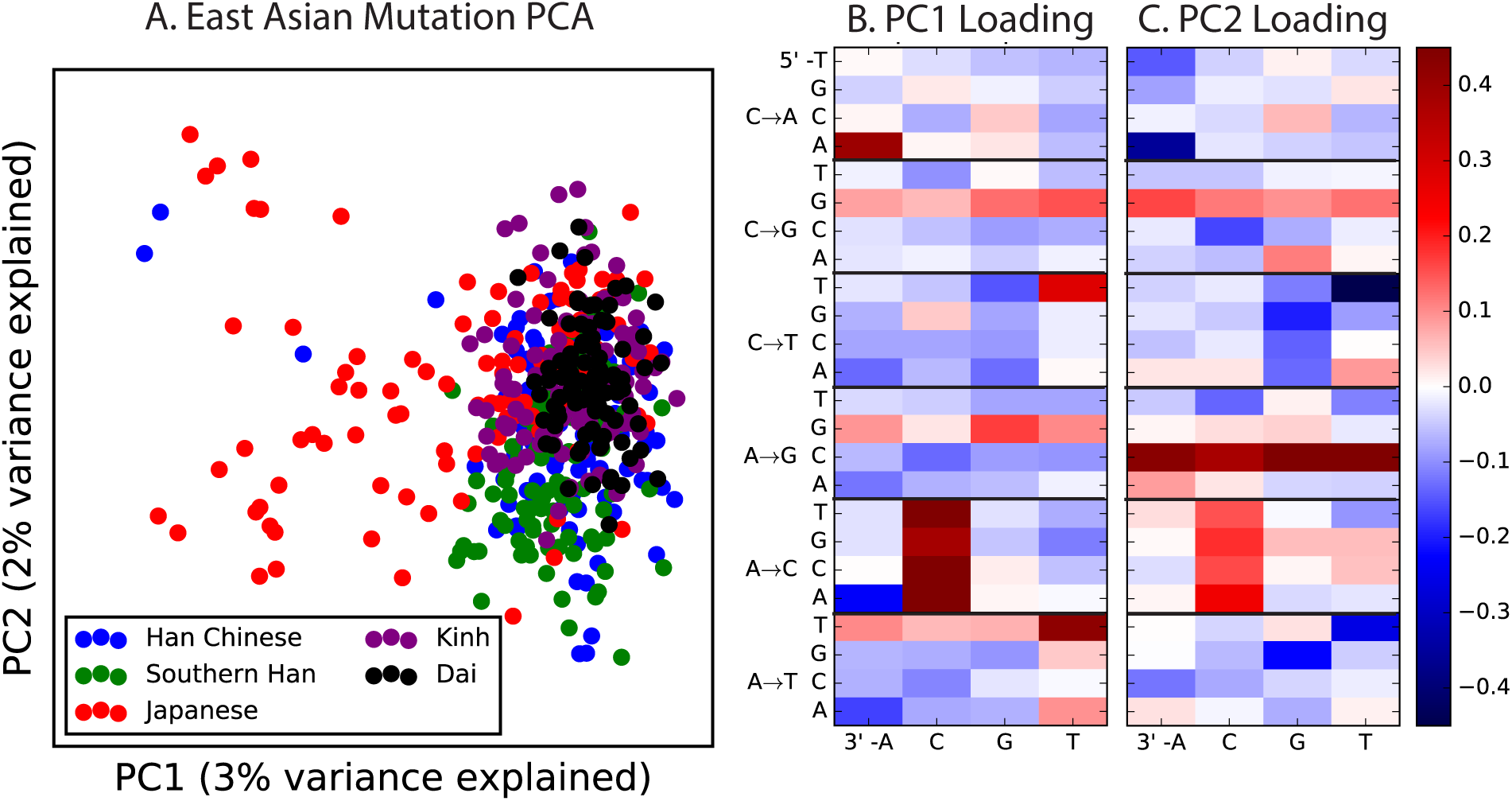
PCA of East Asian populations. Population abbreviations are: CDX = Chinese Dai in Xishuangbanna, China; JPT = Japanese in Tokyo, Japan; CHB = Han Chinese in Bejing, China; CHS = Southern Han Chinese; KHV = Kinh in Ho Chi Minh City, Vietnam.

**Figure 4–Figure Supplement 5:**
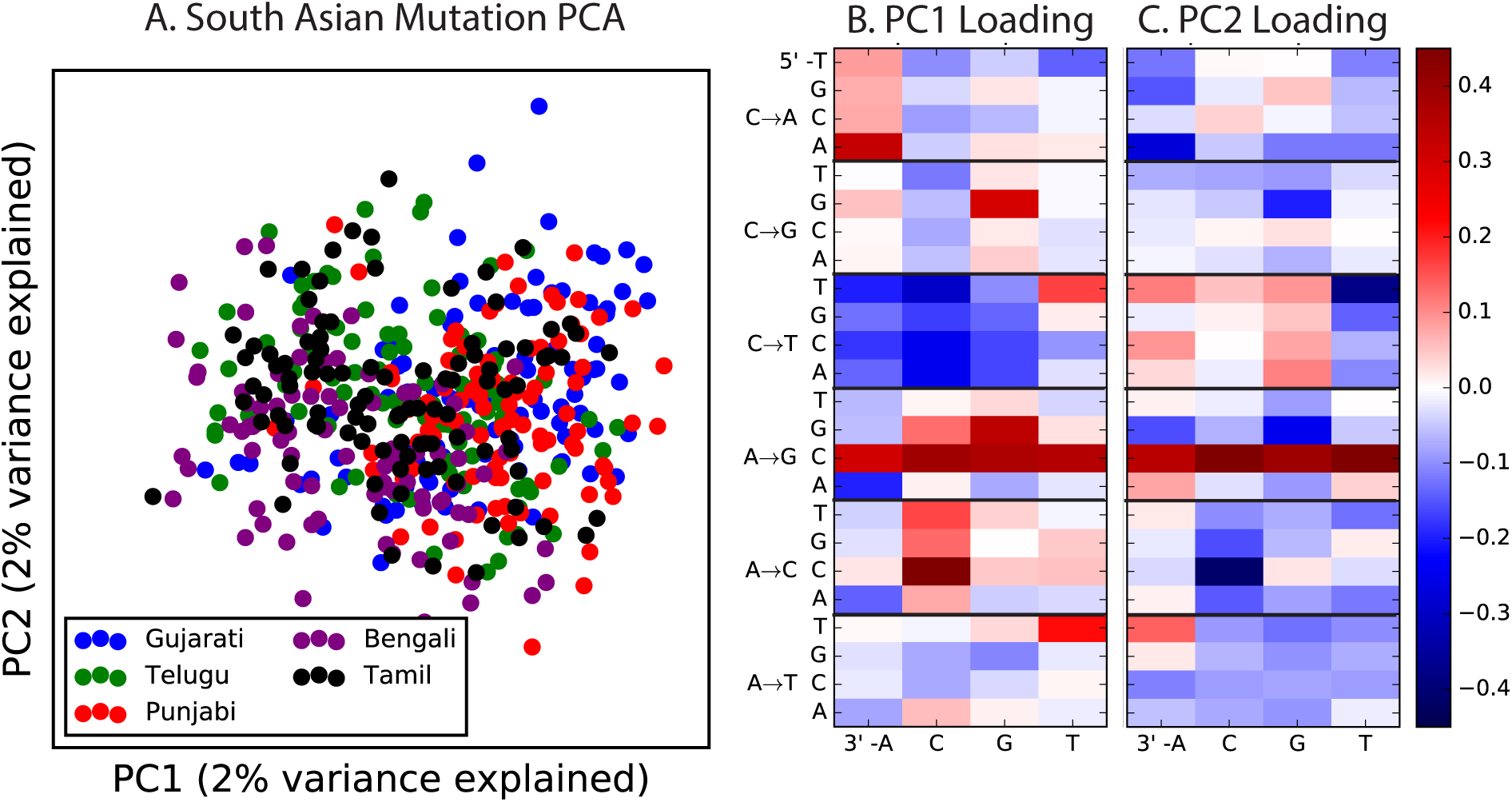
PCA of South Asian populations. Population abbreviations are: ITU = Indian Telugu from the UK; GIH = Gujarati Indian from Houston, Texas; PJL = Punjabi from Lahore, Pakistan; BEB = Bengali from Bangladesh; STU = Sri Lankan Tamil from the UK.

**Figure 4–Figure Supplement 6:**
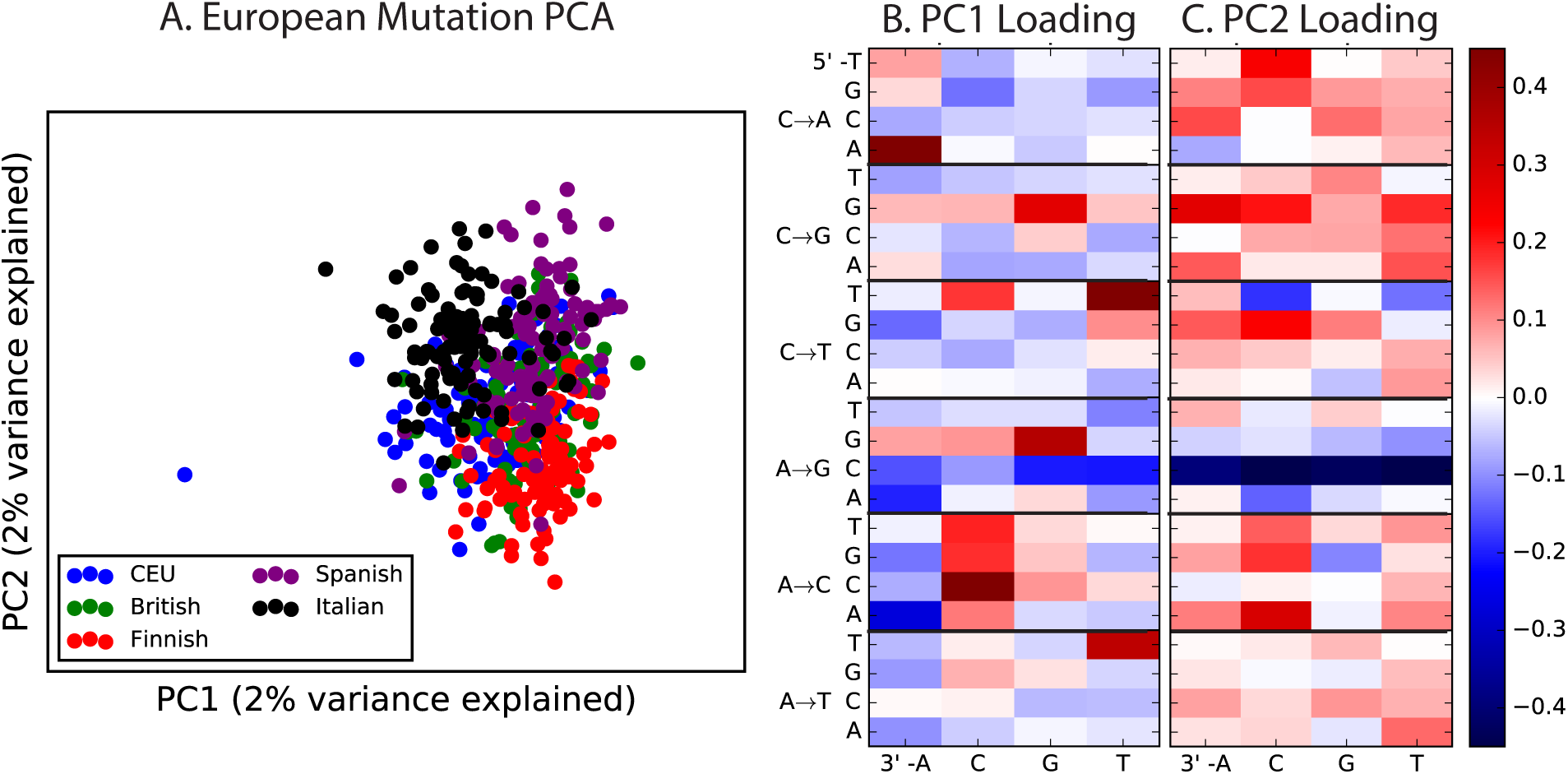
PCA of European populations. Population abbreviations are: IBS = Iberian Population in Spain; TSI = Toscani in Italia; GBR = British in England and Scotland; CEU = Utah Residents (CEPH) with Northern and Western Ancestry; FIN = Finnish in Finland.

**Figure 4–Source Data 1.** This text file shows the number of SNPs in each of the 96 mutational categories that passed all filters in each finescale 1000 Genomes population.

**Figure 5–Figure Supplement 1:**
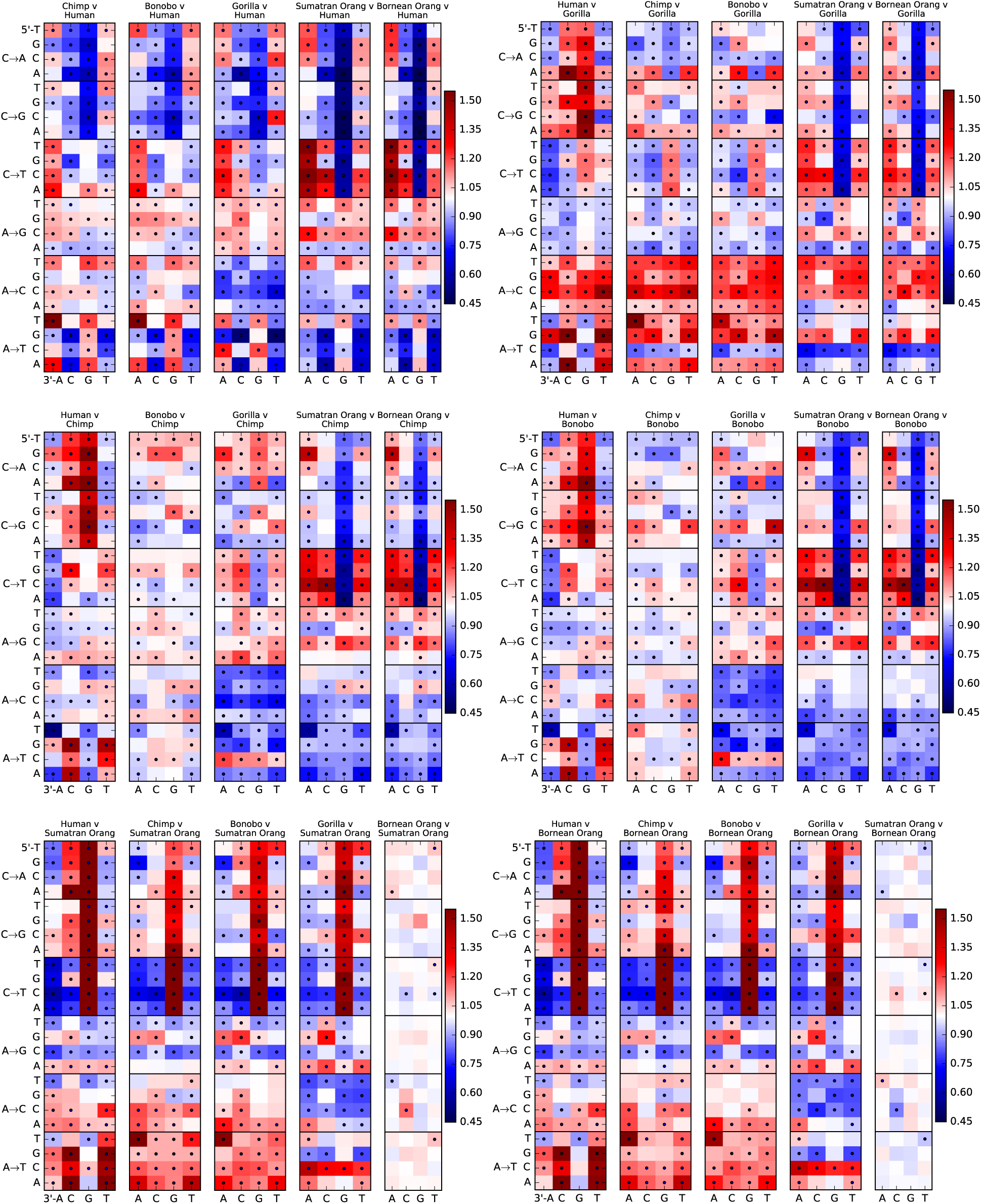
Mutation spectra of great apes. These heatmap comparisons demonstrate that closely related great apes such as Chimpanzees and Bonobos have more similar mutation spectra than more distantly related apes do.

## References

[1] Kong, A. et al. Rate of *de novo* mutations and the importance of father’s age to disease risk. Nature 488, 471–475 (2012).

[2] Ségurel, L., Wyman, M. & Przeworski, M. Determinants of mutation rate variation in the human germline. Annu Rev Genomics Hum Genet 15, 19.1–19.24 (2014).

[3] Alexandrov, L. et al. Signatures of mutational processes in human cancer. Nature 500, 415–421 (2013).

[4] Helleday, T., Eshtad, S. & Nik-Zainal, S. Mechanisms underlying mutational signatures in human cancers. Nature Reviews Genetics 15, 585–598 (2014).

[5] Shinbrot, E. et al. Exonuclease mutations in DNA polymerase epsilon reveal replication strand specific mutation patterns and human origins of replication. Genome research 24, 1740–1750 (2014).

[6] Ohno, M. et al. 8-oxoguanine causes spontaneous *de novo* germline mutations in mice. Scientific Reports 4, 10.1038/srep04689 (2014).

[7] Lynch, M. The lower bound to the evolution of mutation rates. Genome biology and evolution 3, 1107–1118 (2011).

[8] Sung, W., Ackerman, M., Miller, S., Doak, T. & Lynch, M. Drift-barrier hypothesis and mutation-rate evolution. Proc Natl Acad Sci USA 109, 18488–18492 (2012).

[9] Hwang, D. & Green, P. Bayesian Markov chain Monte Carlo sequence analysis reveals varying neutral substitution patterns in mammalian evolution. Proc Natl Acad Sci USA 101, 13994–14001 (2004).

[10] But, D. et al. Mismatch repair incompatibilities in diverse yeast populations. Genetics 205, 10.1534/genetics.116.199513 (2017).

[11] Seoighe, C. & Scally, A. Inference of candidate germline mutator loci in humans from genome-wide haplotype data. PLoS Genetics 13, e1006549 (2017).

[12] Harris, K. Evidence for recent, population-specific evolution of the human mutation rate. Proc Natl Acad Sci USA, 112, 3439–3444 (2015).

[13] Mathieson, I. & Reich, D. E. Variation in mutation rates among human populations. bioRxiv 063578 (2016).

[14] 1000 Genomes Project Consortium et al. A global reference for human genetic variation. Nature 526, 68–74 (2015).

[15] Pollard, K., Hubisz, M., Rosenbloom, K. & Siepel, A. Detection of nonneutral substitution rates on mammalian phylogenies. Genome Research 20, 110–121 (2010).

[16] November, J. et al. Genes mirror geography within Europe. Nature 456, 98–101 (2008).

[17] Mallick, S. et al. The Simons Genome Diversity Project: 300 genomes from 142 diverse populations. Nature 538, 201–206 (2016).

[18] UK10K Consortium. The UK10K project identifies rare variants in health and disease. Nature 526, 82–90 (2015).

[19] Prado-Martinez, J. et al. Great ape genetic diversity and population history. Nature 499, 471–475 (2013).

[20] Moorjani, P., Amorim, C., Arndt, P. & Przeworski, M. Variation in the molecular clock of primates. Proc Natl Acad Sci USA 113, 10607–10612 (2016).

[21] Goodman, M. The role of immunochemical differences in the phyletic development of human behavior. Human Biol 33, 131–162 (1961).

[22] Li, W. & Tanimura, M. The molecular clock runs more slowly in man than in apes and monkeys. Nature 326, 93–96 (1987).

[23] Scally, A. & Durbin, R. Revising the human mutation rate: implications for understanding human evolution. Nature Rev Genetics 13, 745–753 (2012).

[24] Galtier, N., Piganeau, G., Mouchiroud, D. & Duret, L. GC-content evolution in mammalian genomes: the biased gene conversion hypothesis. Genetics 159, 907–911 (2001).

[25] Carlson, J. et al. Extremely rare variants reveal patterns of germline mutation rate heterogeneity in humans. bioRXiv preprint https://doi.org/10.1101/108290 (2017).

[26] Do, R. et al. No evidence that selection has been less effective at removing deleterious mutations in Europeans than in Africans. Nature Genetics 47, 126–131 (2015).

[27] Martin, A. P. & Palumbi, S. R. Body size, metabolic rate, generation time, and the molecular clock. Proceedings of the National Academy of Sciences 90, 4087–4091 (1993).

[28] Amster, G. & Sella, G. Life history effects on the molecular clock of autosomes and 616 sex chromosomes. Proceedings of the National Academy of Sciences 113, 1588–1593 (2016).

[29] Moorjani, P., Gao, Z. & Przeworski, M. Human germline mutation and the erratic molecular clock. bioRxiv 058024 (2016).

[30] Gao, Z., Wyman, M., Sella, G. & Przeworski, M. Interpreting the dependence of mutation rates on age and time. PLoS Biology 14, e1002355 (2016).

[31] Narasimhan, V. et al. A direct multi-generational estate of the human mutation rate 623 from autozygous segments seen in thousands of parentally related individuals. bioRxiv preprint “http://dx.doi.org/10.1101/059436”(206).

[32] Coop, G., Wen, X., Ober, C., Pritchard, J. K. & Przeworski, M. High-resolution mapping of crossovers reveals extensive variation in fine-scale recombination patterns among humans. Science 319, 1395–1398 (2008).

[33] Baudat, F. et al. PRDM9 is a major determinant of meiotic recombination hotspots in humans and mice. Science 327, 836–840 (2010).

[34] Smit, A., Hubley, R. & Green, P. RepeatMasker Open-4.0. http://www.repeatmasker.org (2013-15).

[35] Field, Y. et al. Detection of human adaptation during the past 2,000 years. Science 13, 10.1126/science.aag0776 (2016).

[36] Tennessen, J. et al. Evolution and functional impact of rare coding variation from deep sequencing of human exomes. Science 6, 64–69 (2012).

[37] Hoffman, M. et al. Integrative annotation of regulatory elements from ENCODE data. Nucleic Acids Research 41, 827–841 (2013).

[38] Woodfine, K. et al. Replication timing of the human genome. Hum Mol Genet 13, 191–202 (2004).

[39] Lek, M. et al. Analysis of protein-coding genetic variation in 60,706 humans. Nature 536, 285–291 (2016).

[40] Hernandez, R., Williamson, S. & Bustamante, C. Context dependence, ancestral misidentification, and spurious signatures of natural selection. Mol Biol Evol 24,

